# A wing growth organizer in a hemimetabolous insect suggests wing origin

**DOI:** 10.1101/2021.03.10.434860

**Authors:** Takahiro Ohde, Taro Mito, Teruyuki Niimi

## Abstract

The origin and evolution of insect wings remain enigmatic after a century-long discussion. Molecular dissection of wing development in hemimetabolous insects, in which the first functional wings evolved, is key to understand genetic changes required for wing evolution. We investigated *Drosophila* wing marker genes in the cricket, *Gryllus bimaculatus*, and found *apterous* and *vestigial* show critical functions in nymphal tergal identity and margin formation, respectively. We further demonstrate that margin cells in the lateral-anterior tergal region constitute a growth organizer of wing blades. Transcriptome and RNAi analyses unveiled that Wnt, Fat-Dachsous, and Hippo pathways are involved in disproportional growth of *Gryllus* wings. Our data collectively support the idea that tergal margin cells of a wingless ancestor gave rise to the body wall extension required for evolution of the first powered flight.

## INTRODUCTION

Insects were the first animal group to achieve powered flight. The significance of this development for successful radiation of this animal group is unarguable, yet the evolutionary history of the wing is still unclear after discussion over a century. The dispute has focused primarily on two concepts of wing origin: (1) extension of the dorsal body wall (e.g. Crampton, 1916; Snodgrass, 1935), and (2) modification of a pre-existing structure related to a ventral appendage (e.g. Wigglesworth, 1978; Kukalova-Peck, 1983). These concepts link the wing to anatomically distinct body parts of the insect ground plan, tergum and pleuron, respectively, and evolutionary history of the wing is separately illustrated for each hypothesis (Clark-Hachtel and Tomoyasu, 2016). A dual origin hypothesis reconciles these classical ideas and proposes that both tergal and pleural elements fused to form the first functional wings (Rasnitsyn, 1981; Clark-Hachtel and Tomoyasu, 2016). This idea has gained support in recent developmental and paleontological studies. Still, the hypothesis embraces a spectrum of interpretations on the degree of contribution from each element (Linz and Tomoyasu, 2018). Whereas understanding of the genetic regulation of wing development has accumulated in *Drosophila*, this knowledge is largely lacking in other insects, particularly outside the Holometabola. Holometabolous larvae undergo dynamic remodeling of the entire body organization during pupation to form the adult insect. The evolution of holometabolous development split the life cycle into two modules and facilitated modification of ancestral hemimetablous development (Truman & Riddiford, 2019). This modified mode of development makes it less suitable to directly compare it with the development of apterygote ancestors for understanding wing evolution.

Hemimetabolous insects show no drastic changes in overall body plan between immature nymphs and adults. Body parts that change substantially in size and pattern during the transition include genitalia and wings. In the modern hemimetabolous insects, wings are not visible in early nymph stages, and evaginated wing primordia, called wing pads, progressively expand and become patterned at each molt. The first winged insects developed obviously in hemimetabolous species, and dissecting hemimetabolous wing development at the molecular level is essential for deducing genetic changes responsible for the rise of wings from an ancestral wingless body plan. In the two-spotted cricket, *Gryllus bimaculatus*, wings extend from lateral areas of nymphal thoracic terga (Mashimo and Machida, 2017), but genetic mechanisms underlying this process are not known.

## RESULTS

### *wg, apAB* and *vg* mRNAs expression in developing thoracic terga in *Gryllus* embryos

Given their central roles in the wing development of *Drosophila melanogaster, vestigial* (*vg*), *apterous* (*ap*) and *wingless* (*wg*) were selected as wing marker genes for finding structures homologous to wings in other insects and crustaceans (Averof and Cohen, 1997; Niwa *et al*., 2010; Ohde *et al*., 2013; Clark-Hachtel *et al*., 2013; Hu *et al*., 2019; Clark-Hachtel and Tomoyasu, 2020). *vg* in *Drosophila* embryos is the earliest marker of wing and haltere imaginal disc cells, although its role in determination of imaginal disc fate is unclear (Williams *et al*., 1991). *vg* is required for the selective proliferation and identity formation of the prospective wing region of larval imaginal discs (Williams *et al*., 1991; Kim *et al*., 1996). In late second instar larvae, Ap in the dorsal compartment of the wing disc induces both *vg* boundary enhancer (*vg*BE) activity and *wg* expression at the dorsoventral (DV) boundary via the Notch pathway (Williams *et al*., 1994; Diaz-Benjumea and Cohen, 1995; Kim *et al*., 1995). Subsequently, Wg from DV boundary cells, together with Decapentaplegic (Dpp) from anteroposterior (AP) boundary cells, activates *vg* quadrant enhancer (*vg*QE) to expand the wing compartment in the wing disc (Kim *et al*., 1996; Zecca and Struhl, 2007).

We initially examined expression patterns of *wg, apAB*, and *vg* in stages 6, 7, and 9 embryos of *Gryllus*. Tergal and pleural regions are subdivided during these stages (**Fig. 1A**, Mashimo and Machida, 2017). In the thoracic tergal region, spots of *wg* mRNA expression appear on the posterior side in early stage 7 embryos and become more prominent in late-stage 7 (**Fig. 1B, C**). Additional *wg* expression in lateral to posterior tergal edges is detected in stage 9 (**Fig. 1D**). *wg* is expressed in stripe patterns at the AP boundary of legs in the pleural region, as previously reported (**Fig. 1B**–**D**; Niwa *et al*., 2000). Two *ap* orthologs in the *Gryllus* genome were classified to ApA and ApB from six amino acid residues unique to each group in the homeodomain (**Fig. S1**). However, partial sequences we obtained were insufficient to separately analyze *in situ* expression of each gene (**Fig. S1**). We detected *apAB* signals in thoracic terga of stage 6 embryos that were maintained to stage 9 (**Fig. 1E**–**G**). *apAB* expression covers a broad area of thoracic terga, yet marginal cells lack expression. *apAB* expression is also detected in the thoracic pleural regions in addition to tergal cells. *apAB* is expressed in a U-shape pattern in stage 6, becomes separated into two areas in late stage 7, and forms connected areas, reshaping the U in stage 9 (**Fig. 1E**–**G**). Clear *vg* signals were first detectable at stage 9, in contrast to *wg* and *apAB* (**Fig. 1H**–**J**). Expression localized to tergal margins and two pleural spots (**Fig. 1J**). Marginal expression patterns of *wg* and *vg* in stage 9 embryos are both localized to the lateral to posterior regions of meso- and metathorax (T2 and T3), and in the anterior region of prothorax (T1) (**Fig. 1D, G, J, K**).

**Fig. 1.**
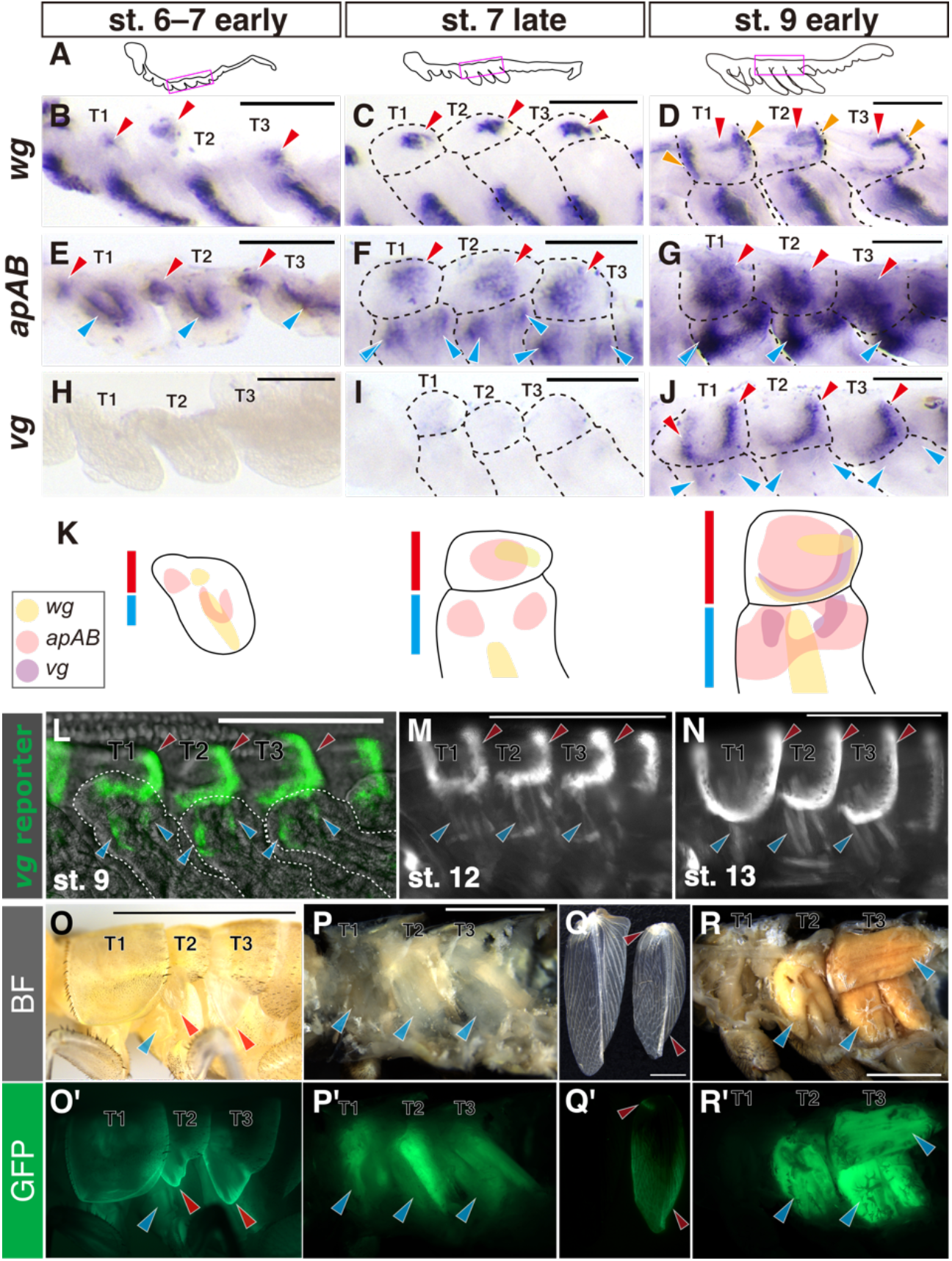
Expression pattern of *wg, apAB* and *vg* in thoracic segments of *Gryllus*. **A**, Schematics illustrate lateral views of *Gryllus* embryos in each stage. Magenta rectangle indicates the region shown in **B**–**J. B–J** Dorsolateral regions of thoracic segments stained for *wg* (**B–D**), *apAB* (**E**–**G**) and *vg* (**H**–**J**) with *in situ* hybridization in stage 6 to early stage 7 (**B, E, H**), late-stage 7 (**C, F, I**) and early-stage 9 (**D, G, J**). **K**, Summary of gene expression pattern in meso- and metathoracic segments. Red and blue lines indicate tergal and pleural regions, respectively. **L**–**N** *vg* reporter gene signals in dorsolateral regions of thoracic segments in late embryos. **O**–**R and O’**–**R’** *vg* reporter gene signals in post-embryonic stages. Dorsolateral region (**O, O’**) and thoracic musculature in median section (**P, P’**) of sixth instar nymph, and forewing (**Q, Q’**) and thoracic musculature in median section (**R, R’**) of adult. Crickets before cuticle coloration are shown for both nymph and adult. Red and blue arrowheads indicate expression in tergal and pleural regions, respectively. Scale bars are 100 µm in **B**–**J**, 250 *µ*m in **L**–**N** and 3 mm in **O**–**R**, respectively.

*vg* is an embryonic marker of wing disc cells in *Drosophila* (Williams *et al*., 1991). The wing disc includes a primordium of the wing and notum and pleural tissues. The formation of this compact imaginal disc is characteristic only in higher Diptera (Nijhout, 1994), and the fate of *vg*-expressing embryonic cells in insects other than *Drosophila* is unknown. In *Gryllus*, the evidence is limited to mRNA expression patterns with *in situ* hybridization in late embryos due to outer cuticle formation that causes strong non-specific staining (Niwa *et al*., 2000). We generated a *vg* reporter line by knocking-in the EGFP cassette at the upstream side of the *vg* coding sequence (**Fig. S2A**–**C**). This reporter line (vg5’GFP) mimics *vg* mRNA expression in both tergal and pleural regions at stage 9 (**Fig. 1K**). During late embryonic development, the EGFP signal pattern in the pleural region is transformed from two spots to several cylindrical shapes (stage 12–), but the tergal region is maintained without significant change from earlier stages (**Fig. 1K**–**M, Fig. S2C**–**Q**). *vg* reporter signal in nymphs appears at the margins of wing pads while strong signaling disappears in the posterior tergal margin regions (**Fig. 1N, N’**). A signal was also detected in thoracic dorsoventral muscles (**Fig. 1O, O’**). *vg* reporter signals were detected in adult wings in a broad area at the distal region and at a spot in the proximal region (**Fig. 1P, P’**). We detected robust reporter activity in adult thoracic musculature including dorsal longitudinal and dorsoventral and pleural muscles that are indirect-and direct flight muscles, respectively (**Fig. 1Q, Q’, Fig. S3**; Furukawa *et al*., 1983).

### *apAB* and *vg* function in nymphal thoracic terga and adult wing formation

We investigated the function of wing marker genes in body plan formation of *Gryllus* by generating CRISPR/Cas9-mediated mosaic knockouts for each gene. A loss of *wg* function likely causes no effects due to functional redundancy (Miyawaki *et al*., 2004), and we focused on *vg* and *apAB* functions. We designed a sgRNA targeting *EGFP* as a negative control (**Fig. 2A**–**E**; **Table S1**). Consistent with the expression pattern during embryogenesis, *vg* sgRNA/Cas9 injected individuals (*vg*^*CRISPR*^) exhibited a lack of tergal margins in both wingless (T1) and wing-bearing (T2 and T3) segments of first instar nymphs (**Fig. 2F**–**H**). Further, individuals without thoracic tergal margins displayed severely reduced wings in adults (**Fig. 2I, J**). Characteristic patterns in thoracic terga in *apAB*^*CRISPR*^ individuals, such as color, shape, and hairs were lost (**Fig. 2K**–**M**). Affected tergal areas lost their normal smooth surface, replaced by the rough surface seen in intersegmental membranes of wildtype insects (**Fig. 2N**). Loss of black color was also induced in lateral-posterior regions of abdominal segments in *apAB*^*CRISPR*^ nymphs (**Fig. 2K**). These nymphs did not survive to adults.

**Fig. 2.**
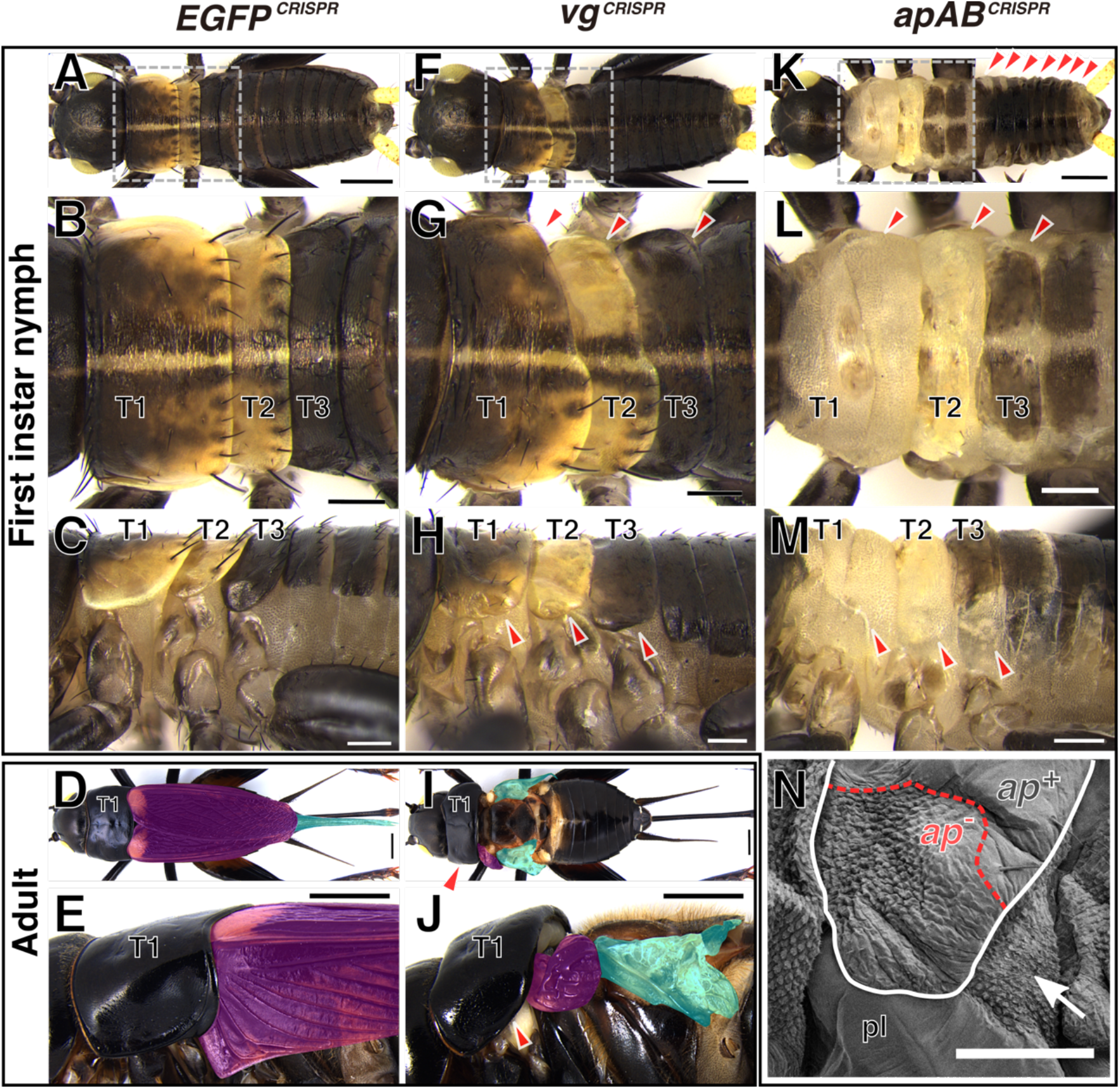
Mosaic knockouts of *vg* and *apAB*. **A**–**M**, Representative images of first instar nymphs (**A**–**C, F**–**H, K**–**M**) and female adults (**D, E, I, J**) injected with Cas9 protein and sgRNAs targeting *EGFP* (**A**–**E**), *vg* (**F**–**J**) and *apAB* (**K**–**M**) in early embryos from dorsal (**A, B, D, F, G, I, K, L**) and lateral (**C, E, H, J, M**) views. Boxed area in **A, F, K** is magnified in **B, G, L**, respectively. Arrowheads indicate regions affected by gene knockouts. Fore- and hindwings are shaded in magenta and cyan, respectively. Arrowheads indicate affected tergal regions. **n** A scanning electron micrograph of the dorsolateral region in mesothorax of an *apAB* mosaic knockout cricket. The tergum is outlined with a white line. Tergal surface with specific (*ap*^-^) and wildtype (*ap*^+^) phenotypes separated by red dotted line. The rough surface structure of *ap*^-^area resembles a soft intersegmental region (arrow). pl, pleuron. Scale bars are 250 *µ*m in **A**–**C, F**–**H, K**–**M**, 2 mm, **D, E, I, J** and 100 µm in **N**.

### The lateral-anterior region of thoracic tergum is essential for wing formation

The severe defect in wing formation that follows the loss of nymphal tergal margins in *vg*^*CRISPR*^ suggests that the tergal margin region associated with *vg* expression during embryogenesis is required for wing growth. During post-embryonic development, lateral regions in T2 and T3 that give rise to wings show dramatic exponential growth, while the rest the same segment exhibits linear growth (**Fig. S4**). *vg* expression and function in *Gryllus* embryos are displayed in lateral and posterior margin regions of T2 and T3, and this growth pattern suggests that the lateral region comprises the wing growth organizer. We ablated a part of the mesothoracic tergum in the third instar nymph to test this hypothesis and analyzed effects on adult wing size (**Fig. 3A**). Adult wing size is unaffected after posterior tergum ablation, while removal of lateral regions severely reduced wing size (**Fig. 3B**). This effect is more prominent when larger regions of the lateral tergum are ablated (**Fig. 3B**). A similar size reduction was found in wing articulation, although the effect is small (**Fig. 3C**).

**Fig. 3.**
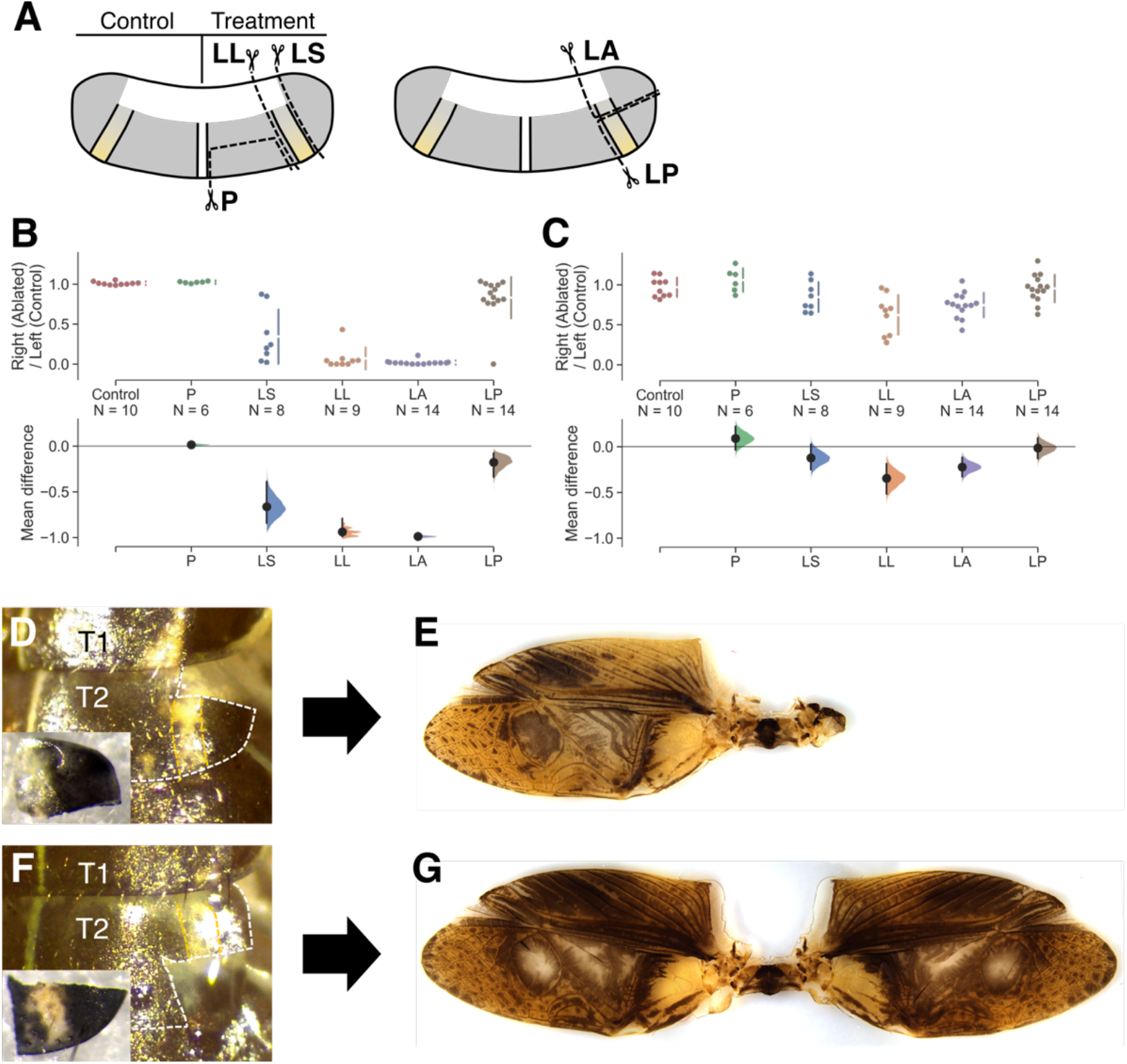
Identification of wing growth organizer in the lateral-anterior region of thoracic terga. **A**, Schematic illustration indicates ablated regions in the third instar nymph: P, posterior; LS, lateral-small; LL, lateral-large; LA, lateral-anterior; LP, lateral-posterior. **B, C**, Effect of nymphal tergal ablations on wing (**B**) and articulation (**C**) sizes in adults. Cumming plots indicate relative area of the treated right side to untreated left side (top) and mean differences of the values from control for wing (**B**) and articulation (**C**) regions (bottom). **D**–**G**, Representative results of the ablation treatment. The right side of targa of the third instar nymph after the ablation of the LA (**D**) and LP (**E**) regions. The ablated tissues are shown in insets. Resultant adult wings of the same individuals are shown in **F** and **G**, respectively. Mesoterga and the yellowish line is outlined with white and yellow dotted lines, respectively.

We independently ablated anterior and posterior regions to further identify the lateral region critical to wing formation (**Fig. 3A**). We found that removal of the anterior region results in almost complete loss of the wing blade, whereas removal of the posterior region has almost no effect on wing size (**Fig. 3B, D**–**G**). Ablation of the lateral-anterior (LA) region of a tergum is sufficient to cause the loss of almost the entire wing blade, but effects are less prominent on the size of the wing articulation (**Fig. 3C**). Thus, the growth organizer of wing blades located in the LA region of terga.

### A combination of transcriptome and RNAi analyses revealed that Wnt, Fat, and Hippo signaling are required for post-embryonic wing growth

We next compared transcriptomes between lateral and central parts of terga (**Fig. 4A**) to identify genes that regulate the growth of lateral tergal regions in T2 and T3. The steroid hormone 20-hydroxyecdysone (20E) promotes wing growth by increasing both cell number and cell size in lepidopteran insects (Nijhout & Grunert, 2002; Nijhout *et al*., 2018). We thus examined the expression level of two ecdysone responsive genes, *E74* and *E75*, as proxies for hemolymph ecdysone titer to determine the timing of tissue sampling for RNA-seq analysis (Ashburner & Richards, 1976; Li *et al*., 2003). *E74* and *E75* in thoracic segments showed expression peaks at 3-and 2.5-days post ecdysis to third instar (DPE), respectively (**Fig. S5**). We selected 3 DPE as a growing stage since total RNA yield is highest among third instar nymphs, suggesting active transcription and protein synthesis (**Fig. S5**).

**Fig. 4.**
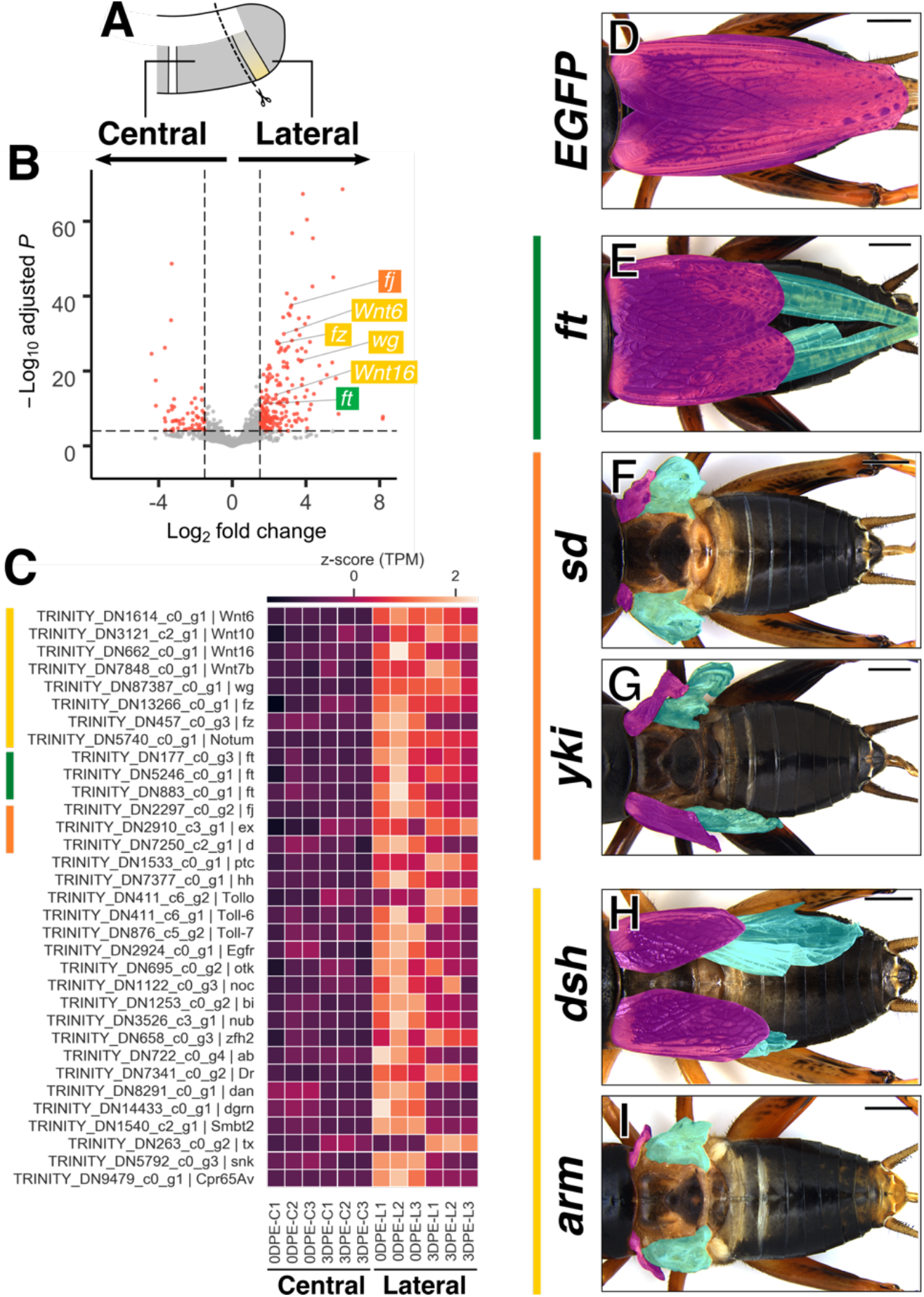
Wnt, Ft-Ds and Hippo signaling are required for the post-embryonic wing growth in *Gryllus*. **A**, Schematics illustrates tergal regions used for the transcriptome comparison. Note that only T2 and T3 segments were used for this analysis. **B**, A volcano plot shows differential expression of transcripts between central and lateral regions in T2 and T3 terga. Data after log-fold change shrinkage with the apeglm method is visualized. Significantly differentially expressed transcripts (adjusted *P* < 10e-5, fold change > 1.5) are in red. Some representative transcripts that are highly expressed in the lateral region are labeled. **C**, Heatmap indicating expression level of 33 candidate genes in each sample. Color code indicates z-score normalized transcripts per million (TPM). **D**–**K**, Adult phenotypes after nymphal RNAi treatment. Fore-and hindwings are shaded in magenta and cyan, respectively. Scale bars are 2 mm.

We obtained raw reads from lateral and central regions of both T2 and T3 in 0 DPE and 3 DPE with a sufficient quality (Q30 >95.87%; **Fig. S6A**). We assembled a transcriptome from these reads with embryonic reads (BUSCO complete orthologs 95.6%; **Fig. S6B, C**), and quantified expression level of each transcript. Transcriptomes display greater similarity within time point than within regions (**Fig. S7A**). We analyzed differentially expressed genes (DEGs) between regions (Lateral versus Central) and between time points (0 DPE versus 3 DPE). We detected more DEGs between time points than between regions, consistent with the above findings (**Fig. S7B**). We presumed 3 DPE as a growing stage, and transcripts highly expressed during this time should be enriched with keywords such as “Developmental protein” and “Cell division”, implying that genes involved in the growth of the lateral region were successfully identified (**Fig. S7C**). DEGs in the lateral region were enriched with the keywords such as “DNA replication”, “Cell cycle”, and “Developmental protein”, support our hypothesis that the lateral terga comprise the growth organizer in T2 and T3 (**Fig. S7C**).

We then proceeded to develop a list of transcripts for RNAi-mediated functional screening. We focused on transcripts annotated as signaling molecules or transcription factors among DEGs more highly expressed in the lateral region than the central region to identify upstream wing growth regulators (**Fig. 4C**). Notably, we found that genes involved in Wnt (Wnt ligands, the receptor, *frizzled*, and the Wnt ligand inactivator, *Notum*), Fat-Ds (the protocadherin, *fat* (*ft*)), and Hippo (the Golgi-kinase, *four-jointed*, the FERM domain gene, *expanded*, and the atypical myosin, *dachs* (*d*)) pathways show high expression in the lateral region. These pathways generate a feed-forward (FF) circuit that plays a central role in the expansion of the wing compartment in *Drosophila* (**Fig. 4B, C**; Zecca & Struhl, 2010). In contrast, components of BMP signaling, another well-characterized signaling pathway that plays a pivotal role in the *Drosophila* wing disc growth and patterning, was not identified as a DEG between regions (**Fig. S8;** Hamaratoglu, *et al*., 2014).

We selected 33 genes, including components of Wnt, Ft-Ds, and Hippo pathways, as candidate growth regulators and performed nymphal RNAi (nRNAi)-mediated functional screening to examine functions in *Gryllus* wing formation (**Fig. 4C, Table S2**). We found specific phenotypes for six transcripts: two *ft* transcripts, *d*, the T-box transcription factor *optomotor-blind* (*omb*), *Epidermal growth factor receptor* (*Egfr*), *zinc finger homeodomain 2* (*zfh2*), and *abrupt* (*ab*). A BLAST-based comparison to the *Drosophila* official gene showed two *Gryllus ft* transcripts that showed effects on the wing size after RNAi treatment derive from a single *ft* locus (**Fig. S9**). Among six genes with specific phenotypes, depletion of *ft* resulted in reduced wing size compared to control *EGFP* dsRNA injected crickets, suggesting a role of Ft-Ds pathways in the *Gryllus* wing growth. *omb* RNAi crickets exhibit irregular wing vein patterns around the AP boundary (**Fig. S10A**–**D**). RNAi treatment of *zfh2* caused high lethality, but surviving adults commonly showed disorganized vein patterns (**Table S2**; **Fig. S10E, F**). *Egfr* RNAi crickets display small adult body sizes, as previously reported (Dabour *et al*., 2011), but did not induce noticeable disproportionate effects on wing size (**Fig. S10G**). We found that *ab* RNAi treatment causes similar small body phenotype (**Fig. S10G**). Depletion of *d*, a regulator of the Hippo pathway, caused irregular male vein patterns and slightly smaller wings although this effect was not clear because of the modest impact (**Fig. S10H**). We targeted genes encoding effectors of Hippo pathway, *scalloped* (*sd*) and *yorkie* (*yki*), to further confirm the role of the Hippo pathway in wing growth. Depletion of *sd* caused a severe reduction in wing size (**Fig. 4H**; **Table S3**). We failed to analyze the effect of *yki* dsRNA injections of third instar nymphs on wing growth due to lethality, but injection of sixth instar nymphs severely reduced wing size (**Fig. 4I**; **Table S3**). In contrast, we found no noticeable effect on wing formation after a single knockdown of each Wnt pathway component (**Table S2**). This result may reflect functional redundancy of ligands and receptors reported in *Gryllus* and other species (Miyawaki *et al*., 2004).

We targeted intercellular components of the canonical Wnt signaling pathway *disheveled* (*dsh*) and *armadillo* (*arm*) to clarify the involvement of the Wnt pathway (**Table S3**). RNAi treatment for both genes resulted in severe reductions in wing size, showing the central role of the Wnt pathway in the *Gryllus* wing growth. Separate RNAi-treatment against *ft, yki, sd, dsh* and *arm* resulted in the reduced size of both female ovipositor and antennae (**Fig. S11A, B**). *d* RNAi crickets displayed curved and short ovipositors (**Fig. S11C**). Ovipositor and antenna size indicate disproportional growth during post-embryonic development, suggesting a shared role of Wnt/Ft-Ds/Hippo pathways in the growth regulation among disproportionally growing organs.

## DISCUSSION

### *vg*-dependent LA cells organize the wing growth in *Gryllus*

In *Gryllus, ap* is expressed across a broad region of terga in stage 9, and *wg* and *vg* show localized expression in thoracic tergal margin cells. Such a spatial gene expression in *Gryllus* embryos is analogous to expression in *Drosophila* wing discs in late second instar larvae. In *Drosophila, ap* induces *vg* and *wg* at the DV boundary to organize wing formation (Kim *et al*., 1995). These three genes show no similar spatial distribution during *Drosophila* embryogenesis. These results demonstrate the timing shift of patterning event from embryonic development in *Gryllus* to post-embryonic development in *Drosophila* (**Fig. 5**). This temporal difference in gene expression might be explained by a developmental modification to form holometabolous larvae. From examples in the development of CNS, legs and eyes, Truman & Riddiford (2019) formulated the principle underlying the evolution of holometabolous larval forms: (1) arrest of ancestral embryonic development programs and (2) redirection of development to an adaptive holometabolous larval form. We ascribe the different timing of the analogous gene expression between *Gryllus* and *Drosophila* to a similar timing shift through the evolution from the hemimetabolous to holometabolous mode. The functional comparison of *ap* and *vg* between the *Drosophila* larval wing disc and the *Gryllus* embryos further supports the concept of a timing shift during evolution, and implicates an ancestral role of *vg* in the formation of the wing organizer. Ap specifies and maintains dorsal cell fate in the *Drosophila* wing disc (Diaz-benjumea & Cohen, 1993). We show *ap* function in the tergal identity formation in the *Gryllus* embryo, indicating an evolutionarily conserved function with different timing of expression between species. *vg* repeatedly plays important roles in *Drosophila* wing development. First, *vg* marks all wing disc-fated cells in embryos by stage 11 (Williams *et al*., 1991). Second, the priming enhancer induces low-level *vg* expression across the entire wing disc. Such expression is required for the initiation of wing growth (Zecca & Struhl, 2008). Third, *vg*BE-regulated *vg* expression organizes feed-forward growth in the wing region (Williams *et al*., 1994; Zecca & Struhl, 2010). Fourth, *vg*QE integrates signals from multiple pathways to expand the wing region (Zecca, 2008; 2010). Analogous expression at the DV boundary between *Drosophila* and *Gryllus* suggests that the third function in organizing wing growth could derive from ancestral hemimetabolous development. Our functional analysis and ablation experiment in *Gryllus* consistently show the lack of a part of the *vg*-dependent tergal margin, the LA region, causes severe wing blade growth defects. In summary, *vg* forms the thoracic tergal margins in the embryonic development, and the *vg*-dependent LA cells organize the wing growth in the postembryonic development.

**Fig. 5.**
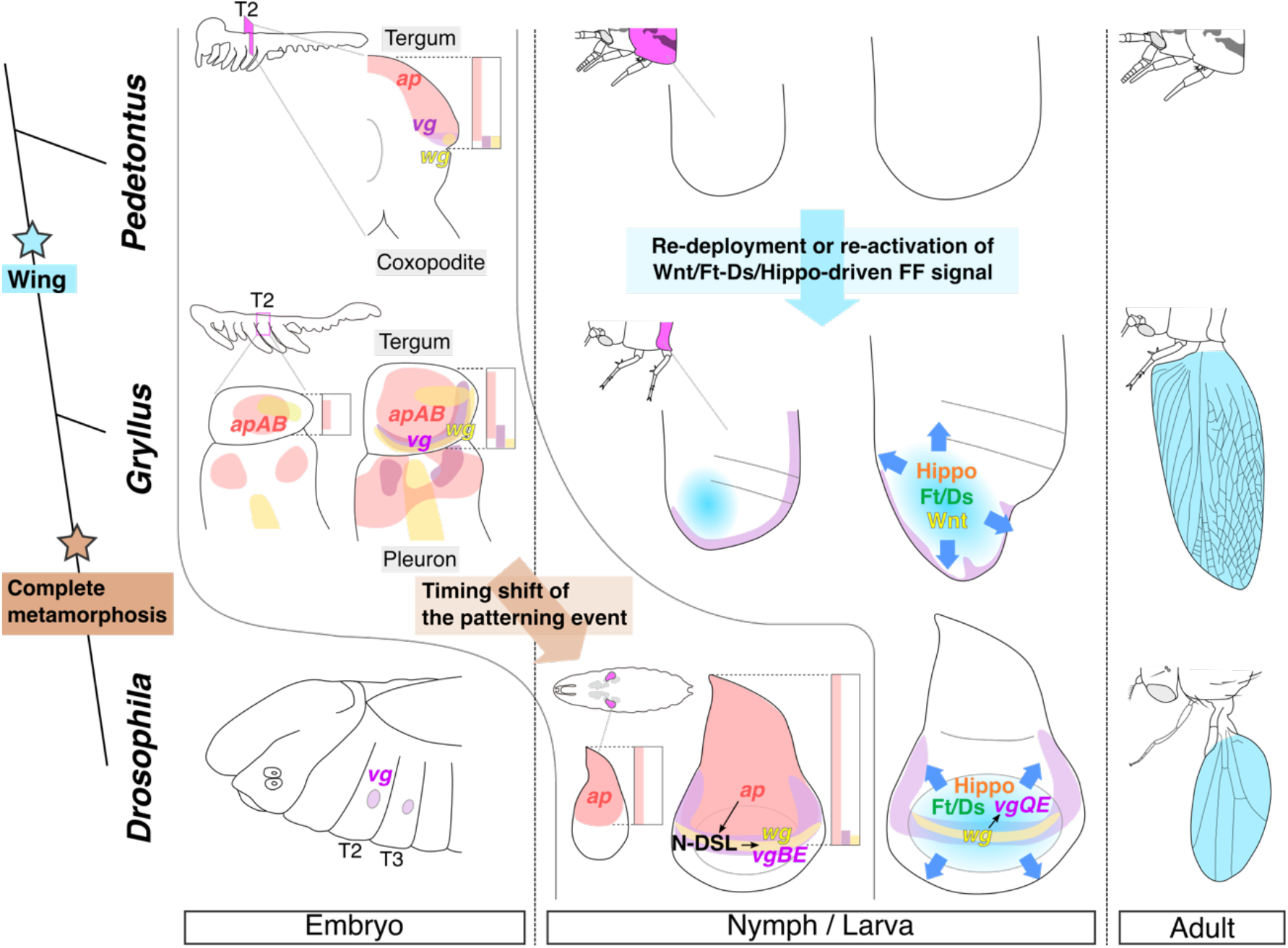
The development and evolution of the wing blade from the tergal margin region. A resemblance of the dorsoventral distribution of *wg*/*ap*/*vg* between *Pedetontus* (Niwa *et al*., 2010) and *Gryllus* indicates that nymphal terga are patterned in a similar fashion. The similar distribution is known in the *Drosophila* wing disc at the late second instar (Kim *et al*., 1995), which implicates a timing shift of the patterning event from embryonic to larval stage. vgBE activity in *Drosophila* is required to initiate expansion of the wing compartment via *vg* FF signaling transduced through actions of vgQE, Wg, and Ft-Ds and Hippo pathways (Zecca and Struhl, 2010). Analogously, the *vg*-dependent tergal margin in *Gryllus* is required for the wing growth, and Wnt, Ft-Ds and Hippo pathways drive dramatic expansion during post-embryonic development. A growth mechanism like that in *Drosophila* was employed at the lateral tergal region of an ancestral apterygote insect to form the body wall extension required for the evolution of insect flight.

### Evolutionarily conserved roles of *vg* and *ap* in flight muscle development

*vg*-expressing cells at the tergal region are critical for the formation of the wing growth organizer, but it is the pleural region that appears to be involved in muscle development. Our detailed reporter analysis showed that the pleural population likely forms myoblasts that differentiate into nymphal dorsoventral muscles. Some nymphal muscles are used for flight in adult insects (Tiegs, 1954). We also detected consistently strong *vg* reporter signals in both indirect and direct flight muscles in T2 and T3 of adults, indicating a likely involvement of *vg* in adult flight muscle development of *Gryllus*. This possibility is concordant with a previous report that *vg* patterns indirect flight muscles, both dorsoventral and dorsal longitudinal, by repressing *ap* and *cut*. These latter genes specify direct flight muscles in *Drosophila* (Bernard *et al*., 2003). We detect *apAB* expression in pleural regions of the *Gryllus* embryo. It suggests evolutionarily conserved functions of *vg* and *ap* in flight muscle development, although the contrasting expression pattern of *vg* in adult direct flight muscles, that is present in *Gryllus* and absent in *Drosophila* (Sudarson *et al*., 2001), implies the modified regulation between species.

### Re-deployment or re-activation of Wnt/Ft-Ds/Hippo pathway-driven FF signals might extend tergal margins of the wingless ancestor

Large wing blades are fundamental to sufficient aerodynamic forces for powered flight. Hence, disproportionate growth of the lateral body wall that gives rise to the wing blade is an essential step for the evolution of functional flight; however, mechanisms underlying this step remain unclear. Each hypothesis for wing origin suggests different scenarios for wing evolution, y*et al*l hypotheses appear to agree on a major contribution of the tergal margin region of the apterygote insect ancestor in the formation of the wing blade (Snodgrass, 1935; Kukalova-Peck, 2008; Prokop *et al*., 2017; Linz and Tomoyasu, 2018; Clark-Hacktel *et al*., 2021; Tomoyasu, 2021). The location of a wing growth organizer at the lateral tergal region in *Gryllus* is concordant with this view. Distribution of *wg, ap* and *vg* mRNAs along the DV axis in *Gryllus* is similar to the distribution in embryos of apterygote bristletail, *Pedetontus unimaculatus* (**Fig. 5**; Niwa *et al*., 2010). Thoracic lateral tergal regions between apterygote and pterygote insects are thus homologous, and some changes in homologous cells led to body wall extension.

Our functional analysis suggests that some changes in Wnt/Ft-Ds/Hippo pathways drove dramatic growth of the homologous region to achieve powered flight. We showed critical roles of Wnt/Ft-Ds/Hippo pathways in the post-embryonic wing growth of *Gryllus*. Together with *vg*, these pathways comprise an FF signal circuit to expand the wing region in the *Drosophila* imaginal disc (Zecca and Struhl, 2010). *Oncopeltus fasciatus*, another hemimetabolous insect, exhibits repression of wing growth after post-embryonic depletion of *vg* (Medved *et al*., 2015). These data suggest that the core wing growth mechanism is conserved between *Gryllus* and *Drosophila* (**Fig. 5**). Intriguingly, recent developmental studies revealed localized expression and function of *vg, sd* and *wg* in crustacean flat epidermal outgrowths such as carapace, tergal edge and coxal plate, although homology of these flat outgrowths to insect wings remains controversial (Shiga *et al*., 2017; Clark-Hachtel and Tomoyasu, 2020; Bruce and Patel, 2020). Because these three genes are components of *vg* FF signal, it suggests crustacean origin of this modular growth and either redeployment or reactivation in the lateral tergal cells extends the body wall of the wingless ancestor that led to insect wings.

## MATERIALS AND METHODS

### Animal husbandry

*Gryllus bimaculatus* strain used in this study derives from the *gwhite* strain (Niwa *et al*., 1997). Cricket colonies were reared in an air-conditioned room at 28–30 °C with 12L:12D photoperiod, and fed on artificial fish food (Spectrum Brands) and cat food (Purina One, Nestlé Purina Petcare). We kept nymphs for staging in an incubator (Panasonic) at 29 °C to minimize developmental stage variation at a time point. Embryo staging followed Donoughe and Extavour (2015).

### *In situ* hybridization

We searched *vg* and *ap* orthologs from *Gryllus* draft genome with BLAST and identified partial nucleotide sequences (*vg*: LC589559, *apA*: LC589561 and *apB*: LC589562; Ylla *et al*., 2021). The public sequence was used for *wg* (AB044713.1). Digoxigenin-labeled riboprobes were transcribed *in vitro* from 188 to 523 bp DNA fragments amplified with primers that have either a SP6 or T7 promoter sequence at the 5′ end (**Table S4**). Embryos were dissected in PBS, and fixed in 4% formaldehyde overnight at 4°C. Fixed embryos were washed in PTx (PBS, 0.1% TritonX-100), dehydrated in a MeOH series, and stored in 100% MeOH at -30 °C until use.

The following procedures were performed at room temperature otherwise described. Fixed embryos were rehydrated in a MeOH series, digested in 2 µg/ml Proteinase K for 5 min, and postfixed in 4% formaldehyde for 20 min. After prehybridization in hybridization buffer (50% deionized formamide, 5X SSC, 100 µg/ml heparin, 100 µg/ml yeast RNA, 0.1% TritonX-100, 0.1% CHAPS, 2% Roche blocking reagent) for more than an hour at 60–65 °C, 200–500 ng of a riboprobe were hybridized with gentle shaking for more than 60 hours at 60–65 °C. Hybridized embryos were washed with a series of SSC buffer at the hybridized temperature, then with maleic acid buffer, and then blocked in 1.5% Roche blocking reagent (11096176001, Merck & Co.) for more than an hour. Blocked embryos were incubated in 1:2,000 anti-Digoxigenin-AP (11093274910, Merck & Co.) overnight at 4°C. After several washes with 1.5% Roche blocking reagent and maleic acid buffer, color was developed with NBT/BCIP solution.

### Size measurement

We collected all samples from a single colony for minimizing variation from genetic and environmental background for assessment of area measurement in central and lateral regions in thoracic segments. We kept all hatched first instar nymphs in a plastic cage, and collect six or seven individuals after each molting. Each sex was separately collected from the fourth instar nymph to the adult. Nymphs and adults were digested in lactic acid overnight at 65°C after cutting off unnecessary body parts, and remaining tergal cuticles of thoracic segments were mounted on glass slides. We mounted nymph specimens in Hoyer’s media. Adult specimens were washed in 100% EtOH and mounted in EUKITT neo (O. Kindler ORSAtec).

Images of mounted slides were captured with a DFC7000 T equipped with a M165 FC (Leica Microsystemsy) and analyzed with ImageJ (version 2.0.0).

### CRISPR/Cas9-mediated genome editing

We assembled oligonucleotides and transcribed and purified single-guide (sg) RNAs with a precision gRNA synthesis kit (A29377, ThermoFisher Scientific,) according to the manufacturer’s instruction. Oligonucleotides used for DNA template assembly are shown in **Table S4**. Synthesized sgRNAs were aliquoted in small volumes and stored at -80°C until use. For microinjection, eggs laid within 1–2 hours were collected from wet paper towels, and pieces of cotton layered in plastic dishes and were aligned in wells in a 2% agarose gel after a brief wash in tap water. A pulled glass capillary connected to a Femtojet microinjector (Eppendolf) was used to inject small droplets of solution into eggs. Materials were injected within 4 hours after egg oviposition.

For somatic gene knockouts, we injected 100 ng/µl sgRNA designed for each gene, and 500 ng/µl *Streptococcus pyogenes* Cas9 protein (Integrated DNA Technologies). Non-homologous end joining (NHEJ)-mediated gene knock-in is performed as previously described for generating the *vg* reporter line (Watanabe *et al*., 2017). The donor plasmid was generated by integrating a partial DsRed sequence as sgRNA target and EGFP expression cassette driven by *Gryllus* cytoplasmic actin promoter (Zhang *et al*., 2002). A sgRNA targeting 5′ region of the *vg* protein-coding sequence was designed with a partial sequence obtained by BLAST search against the *Gryllus* genome assembly (accession no. LC589560). We injected a solution containing 40 ng/µl of sgRNAs targeting the *vg* upstream site and the donor plasmid, 100 ng/µl of the donor plasmid, and 100 ng/µl of Cas9 mRNA transcribed from the linearized MLM3613 plasmid (Addgene 42251; Dahlem *et al*., 2012). GFP-positive individuals were crossed to wildtype, and individuals with GFP signal were selected for establishing the *vg* reporter line.

### Tergum ablation

The ablation was performed on third instar nymphs within 24 hours of molting. Nymphs were anesthetized on ice, and a targeted region of mesotergum on the right-hand side was ablated with a pair of spring scissors. The yellowish line is used as a landmark for ablating different sizes of tissue (i.e. LS and LL in **Fig. 3A**). Ablated nymphs were separately kept in a plastic cup until eclosion, and mesoterga of adults were digested with lactic acid and mounted on glass slides for area measurement. Cumming plots were created with DABEST (Ho *et al*., 2019).

### Quantitative PCR (qPCR)

Nymphs newly molted within two hours to the third instar were periodically collected from a colony and separately kept in small plastic cups at a 29°C. Three to four nymphs were moved on ice for anesthetization at each time point, and thoracic segments without an alimentary canal were dissected in ice-cold PBS, collected individually in TRIzol (Thermo Fisher Scientific), and stored at -80 °C until use. Total RNA was extracted according to manufacturer’s instructions, RNA pellets were resuspended in 30 µl of distilled water and quantified with a Nanodrop 2000 spectrophotometer (Thermo Fisher Scientific). We used 500 ng of total RNA for cDNA synthesis using ReverTra Ace qPCR RT Master Mix with gDNA Remover (Toyobo), and one µl of ten-fold diluted cDNA in ten µl qPCR reaction with THUNDERBIRD SYBR qPCR Mix (Toyobo). A Thermal Cycler Dice Real-Time System II (Takara Bio) was used for qPCR reaction, and we calculated relative gene expression level with the delta Ct method.

### mRNA-sequencing analysis

Nymphs molted within three hours to the third instar were periodically collected from a colony and separately kept in small plastic cups at a 29°C incubator. Zero and 3 DPE samples were anesthetized on ice within 10 min to 2.5 hours and 69 to 73 hours post ecdysis, respectively. Lateral and central tergal parts of both meso- and metathorax were dissected on a paper towel on ice, and then washed in PBS. Dissected tissues from 25 nymphs for a sample were collected in TRIzol (Thermo Fisher Scientific) and stored at -80°C until use. Total RNA was extracted following the manufacturer’s instructions and was further purified with a RNeasy MinElute spin column (QIAGEN). We also extracted total RNA from embryos incubated at 29°C at days 2.5, 3, 3.5, 4.5, 5.5 and 6 after egg laying. Total RNA from terga and embryos were eluted from a column with 20 µl of RNase-free water and submitted to GENEWIZ and Filgen standard RNA-seq service (GENEWIZ) that provided a standard Illumina mRNA library and generated 150 bp pair-end reads on an Illumina HiSeqX and HiSeq4000 (Illumina), respectively. Raw reads were deposited to DDBJ/EBI/NCBI database (PRJDB10701).

We used trimmed reads longer than 25 bp after adaptor sequence removal by Cutadapt (v2.9; Martin, 2011) for the following sequence analysis. We assembled all reads from both tergum and embryo samples together with Trinity (v2.8.4; Grabherr *et al*., 2011), and assessed quality of assembly with BUSCO (v4.0.5; Seppey *et al*., 2019). After quantification of the expression level of each transcript with Salmon (v1.0.1; Patro *et al*., 2017), DEGs were statistically identified with DESeq2 (v1.26.0; Love *et al*., 2014). Functional annotation of assembled transcripts was performed with both Trinotate (v3.1.0) and BLASTX (v2.9.0) against the latest *Drosophila* protein sequences (r6.32). The FlyBase ID of the best BLAST hits in the search against *Drosophila* database was assigned to each transcript and used for gene set analysis with DAVID (v6.8; Dennis *et al*., 2003). Data were visualized with EnhancedVolcano (v1.7.10; Blighe *et al*., 2020), Python3 (v3.7.7) and R (v3.6.3).

### Nymphal RNAi

For the functional screening, we used a consensus sequence created from all isoforms for a target gene in the transcriptome to design a double-stranded RNA (dsRNA). The specificity of dsRNA was assessed by BLAST search to both the transcriptome assembled in this study and the genome (Ylla *et al*., 2021).

RNA was transcribed *in vitro* from a PCR product as a template. PCR primers are listed in **Table S4**. Transcribed RNA was digested with DNase I, then purified with a standard phenol/chloroform extraction protocol. After annealing, dsRNA was aliquoted and stored at -80 °C until use. One µl of dsRNA was injected into the ventral side of the intersegmental membrane between second and third thoracic segments with a pulled glass capillary. Nymphs injected with the same dsRNA were kept in a plastic cage until reaching adulthood.

### Image analysis

Epifluorescent/Confocal/scanning electron microscopy images were taken with an M165 FC (Leica microsystems)/A1R MP (Nikon)/VHX-D500 (KEYENCE), respectively. Images were processed with either ImageJ (version 2.0.0) or GIMP (version 2.10) and assembled and annotated with Inkscape (version 1.0 beta).

## Acknowledgements

We thank Drs. Tetsuya Bando and Sumihare Noji for their help in preliminary analysis of *Gryllus vg* and *sd* function. We thank Dr. Toshiya Ando, and Dr. Takaaki Daimon and his lab members for helpful discussion. We also thank Drs. Yuji Matsuoka, Takahito Watanabe, Yohei Katoh, Taro Nakamura, Shinichi Morita, Hajime Ono, Miki Sugimoto, and Mr. Takahisa Yamashita for technical supports. The computational resource for the RNA-seq analysis was provided by NIG supercomputer system. This study was supported by MEXT KAKENHI (16K18825 and 19H02970 for TO, 16H02596 for TN).

