## Supplemental Data for "A wing growth organizer in a hemimetabolous insect suggests wing origin"

**This PDF file includes:**

Figs S1 to S11

Tables S1 to S4

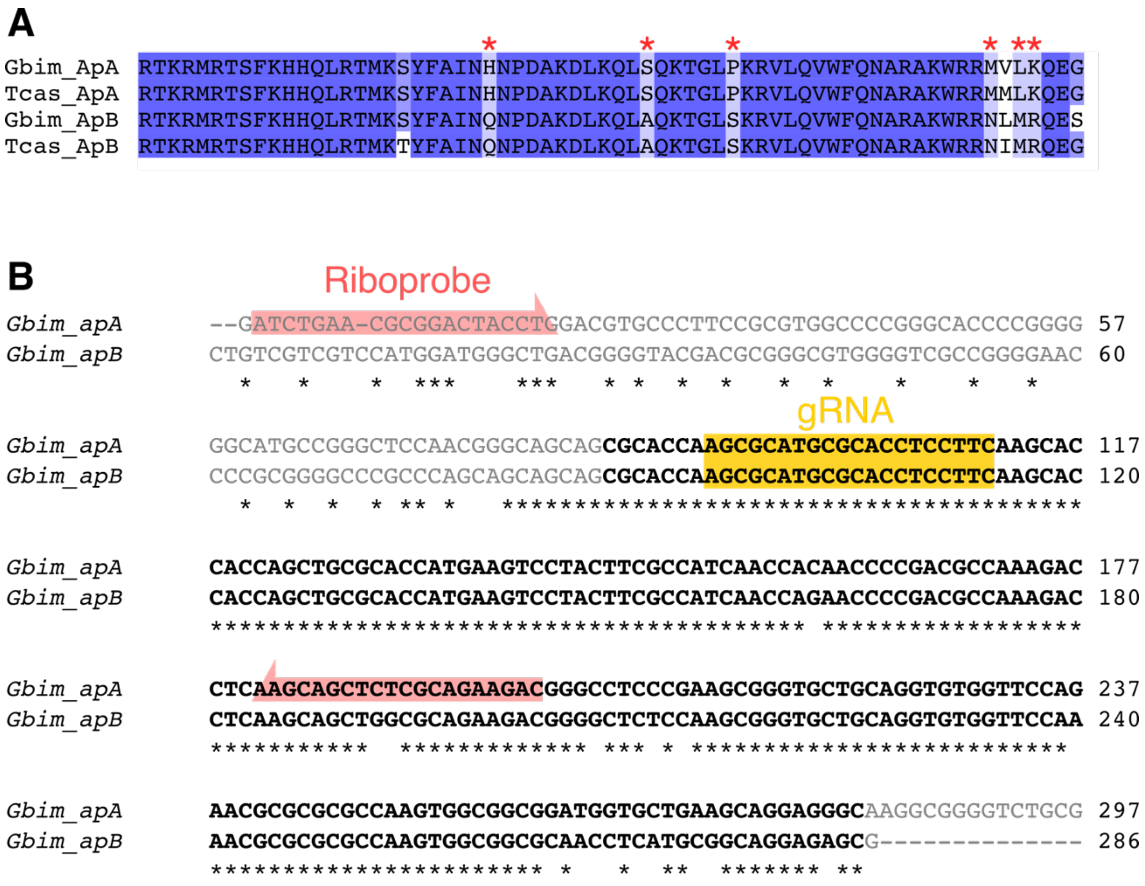

**Fig. S1 *ap* paralogs in *Gryllus***

**A**, The multiple alignment of the Ap homeodomains from *Gryllus bimaculatus* (Gbim) and *Tribolium castaneum* (Tcas). Asterisks indicate amino acid residues unique to either ApA or ApB. **B**, A partial pairwise alignment of *apA* and *apB* identified in this study. Asterisks indicate matched nucleotides. Bold black text indicates homeodomain sequences. The half arrow and boxes in different colors indicate sequences of PCR primers for synthesizing the riboprobe and design of gRNA, respectively.

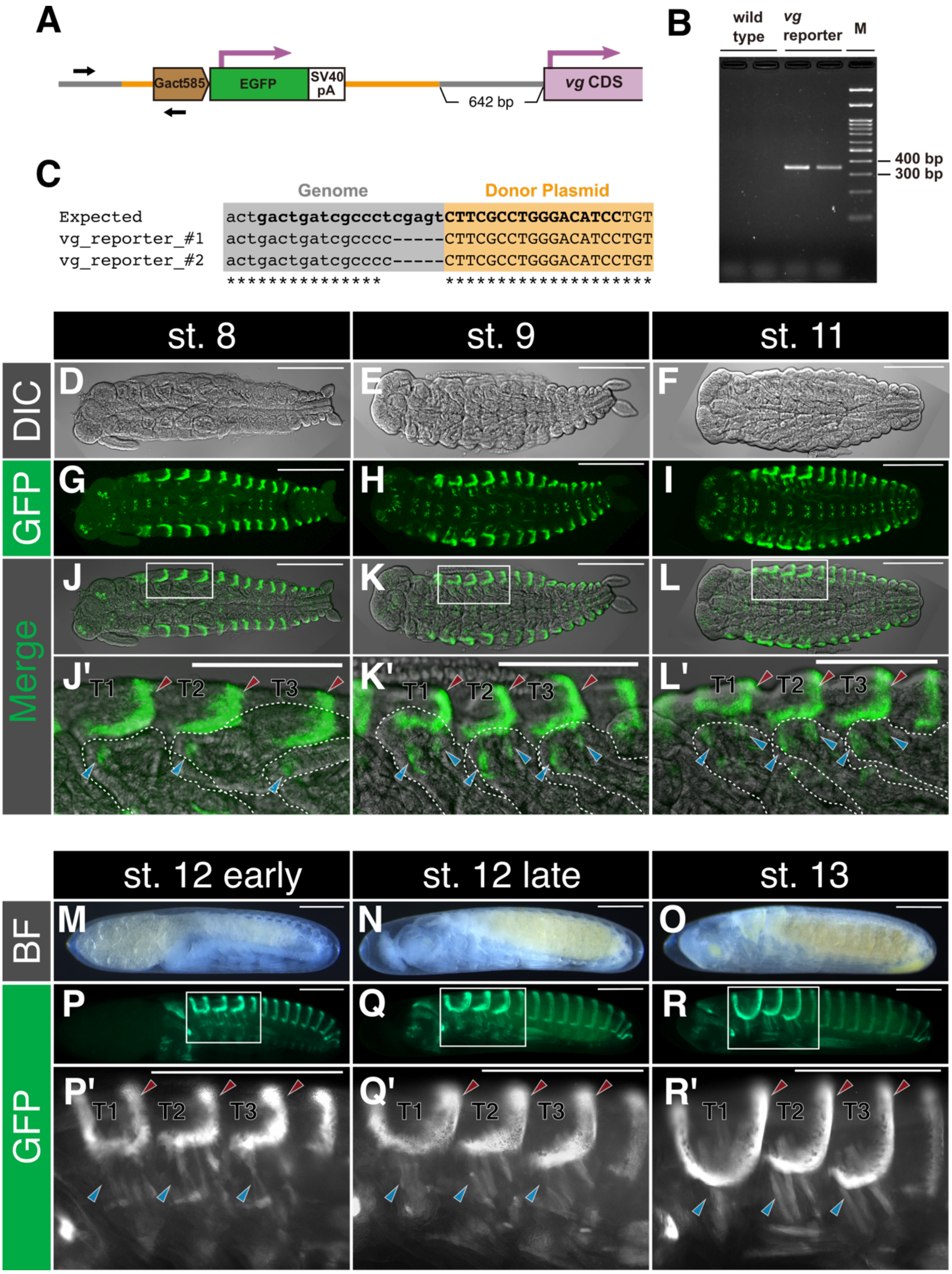
A
Gact585
EGFP
SV40 pA
642 bp
vg CDS
B
wild type
vg reporter
M
400 bp
300 bp
C
Genome
Donor Plasmid
Expected
vg_reporter_#1
vg_reporter_#2
actgactgatcgccctcgagtCTTCGCCTGGGACATCCTGT
actgactgatcgcccc-----CTTCGCCTGGGACATCCTGT
actgactgatcgcccc-----CTTCGCCTGGGACATCCTGT
st. 8
st. 9
st. 11
DIC
GFP
Merge
D
E
F
G
H
I
J
K
L
J'
K'
L'
T1
T2
T3
st. 12 early
st. 12 late
st. 13
BF
M
N
O
P
Q
R
P'
Q'
R'

**Fig. S2 *vg* reporter gene expression during embryogenesis**

**A**, Schematic illustration of the upstream *vg* coding sequence (CDS) with the EGFP reporter cassette. Gray and orange indicate genome and donor plasmid sequences, respectively. Line length is not scaled to sequence length. Black arrows indicate the PCR primer pair used for confirming the insertion. **B**, A gel image showing genomic PCR products in the *vg* reporter line. **C**, DNA sequences around the boundary between the *Gryllus* genome (lowercase) and the inserted donor plasmid (uppercase). “Expected” shows the expected sequence when genome editing was designed. Bold texts indicate sequences targeted by gRNAs. **D**–**L**, Confocal images of vg5´GFP embryos. Maximum projections of confocal stacks are displayed. Differential interference contrast (DIC) microscopy images (**D**-**F**) overlaid with GFP signals (**G**–**I**) are shown in **J**–**L**. **M**–**R** Epifluorescence microscopy images of vg5´GFP embryos from a lateral view. Brightfield (BF) images (**M**–**O**) and GFP signals (**P**–**R**) are shown. Boxed areas in **J**–**L** and **P**–**R** are magnified in **J’**–**L’** and **P’**–**R’**, respectively. Scale bars are 500 $\mu$m and 250 $\mu$m in **D**–**R** and **P’**–**R’**, and **J’**–**L’**, respectively.

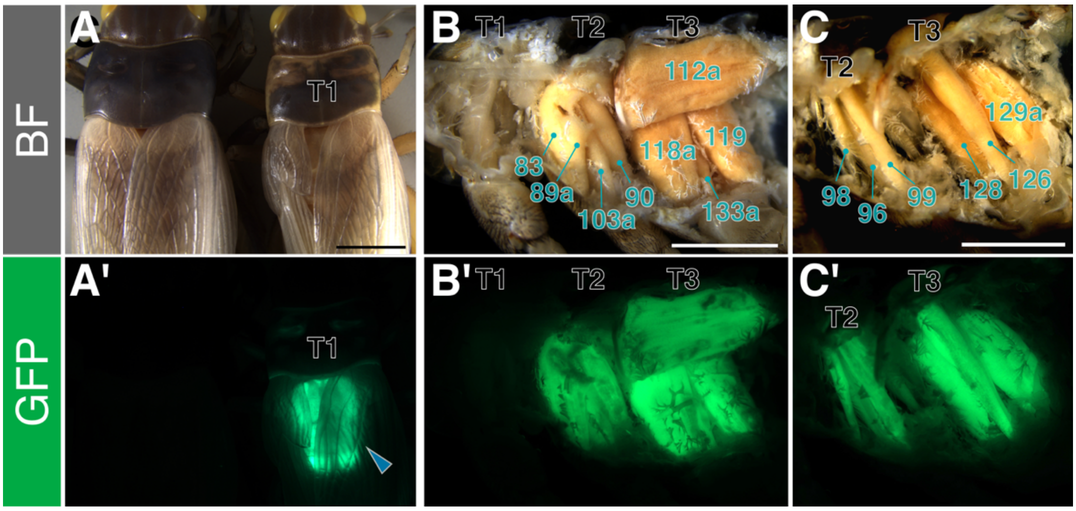

**Fig. S3 *vg* reporter gene expression in adult stage**

**A**, **A'**, Dorsal view of vg5′GFP strain (right) compared to wildtype (left). Crickets before cuticle coloration are shown. Arrowhead indicates GFP signal in thoracic muscles. **B**, **B'**, Dorsal longitudinal muscle (112a) and dorsoventral muscles (the other numbered muscles) in median section. **C**, **C'**, Pleural muscles (all numbered muscles) after removal of medial muscles. Only GFP-positive muscles are annotated, according to Furukawa *et al*. (1983). Scale bar is 3 mm.

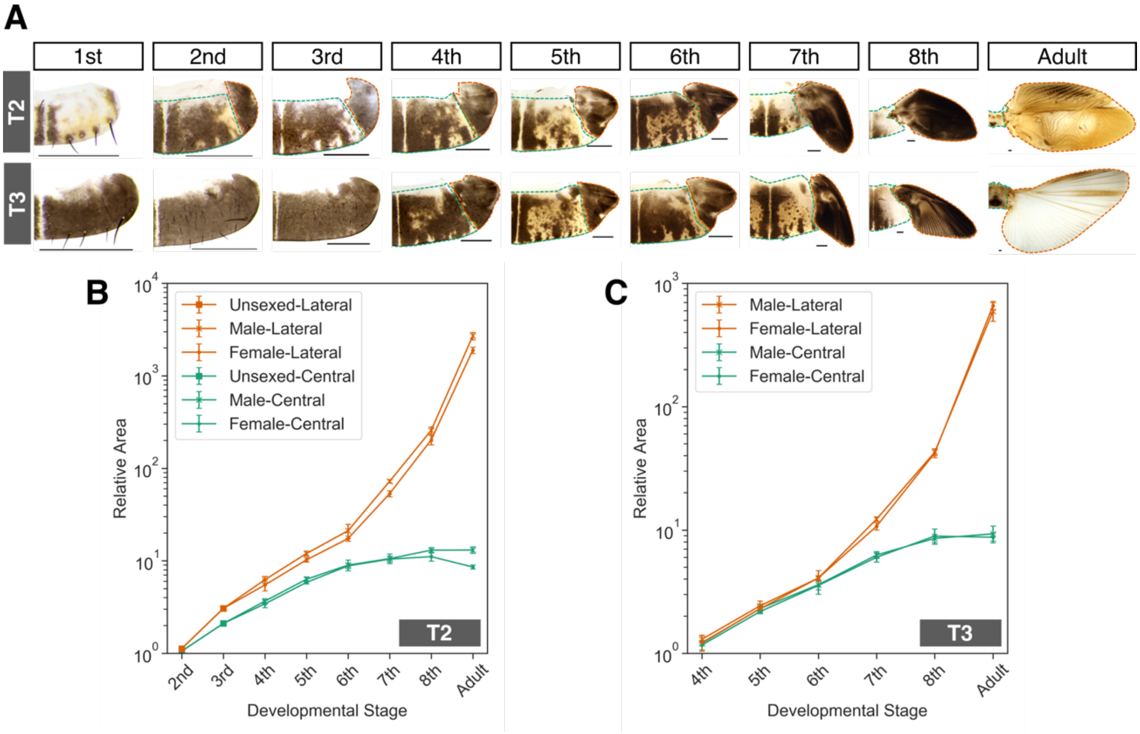

**Fig. S4 Exponential growth of lateral terga**

**A**, Representative image of right halves of terga in mesothorax (T2) and metathorax (T3) from first instar nymph to adult. Unsexed and male terga from first to third instar nymphs and from fourth instar nymph to adult are displayed, respectively. Anterior is up. Orange and green dashed lines outline the lateral and central areas, respectively. Scale bar is 0.5 mm. **B**, **C**, Size of lateral and central regions in T2 (**B**) and T3 (**C**) during post-embryonic development. The smallest area in each data series is set to 1, and relative areas are plotted for the rest. Mean and standard deviation are shown.

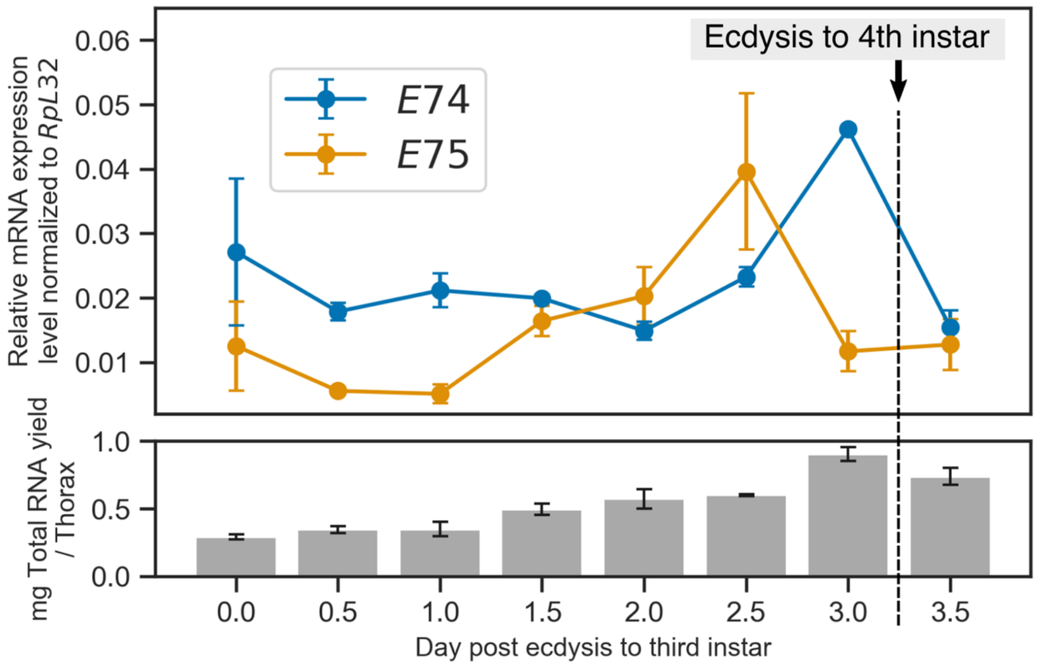

**Fig. S5 Expression level of *E74* and *E75*, and total RNA abundances during third instar nymph**

Mean and standard deviation are shown.

**A**

| SampleID | No. of raw read | Q30(%) | Reads after cutadapt filtering (%reads after cutadapt / raw reads) | Summary of quantification with Salmon | | |
|---|---|---|---|---|---|---|
| | | | | num_processed | num_mapped | percent_mapped |
| d0L1 | 23,432,749 | 96.38 | 23,288,008 (99.4) | 23,288,008 | 22,695,406 | 97.46 |
| d0L2 | 23,467,516 | 96.09 | 23,374,833 (99.6) | 23,374,833 | 22,626,098 | 96.80 |
| d0L3 | 26,132,970 | 96.26 | 26,019,161 (99.6) | 26,019,161 | 25,088,571 | 96.42 |
| d0C1 | 25,329,751 | 96.41 | 25,192,068 (99.5) | 25,192,068 | 24,583,786 | 97.59 |
| d0C2 | 19,455,834 | 96.27 | 19,372,273 (99.6) | 19,372,273 | 18,834,042 | 97.22 |
| d0C3 | 19,568,996 | 96.44 | 19,499,781 (99.6) | 19,499,781 | 18,967,121 | 97.27 |
| d3L1 | 20,301,543 | 96.01 | 20,234,065 (99.7) | 20,234,065 | 19,682,681 | 97.27 |
| d3L2 | 27,346,903 | 95.87 | 27,247,800 (99.6) | 27,247,800 | 26,435,127 | 97.02 |
| d3L3 | 21,023,008 | 96.03 | 20,936,611 (99.6) | 20,936,611 | 20,361,017 | 97.25 |
| d3C1 | 25,952,754 | 96.18 | 25,890,802 (99.8) | 25,890,802 | 25,177,544 | 97.25 |
| d3C2 | 21,088,923 | 96.06 | 20,977,352 (99.5) | 20,977,352 | 20,434,356 | 97.41 |
| d3C3 | 25,111,120 | 96.15 | 24,972,627 (99.4) | 24,972,627 | 24,293,625 | 97.28 |

**B**

| | |
|---|---|
| Total trinity 'genes' | 277,660 |
| Total trinity transcripts | 356,490 |
| Contig N50 (All) | 3,126 |
| Contig N50 (longest) | 874 |
| Total assembled base (all) | 368,966,335 |
| Total assembled base (longest) | 170,990,352 |

**C**

| | |
|---|---|
| n | 1,367 |
| Complete and single-copy (%) | 31.5 |
| Complete and duplicated (%) | 64.1 |
| Fragmented (%) | 0.8 |
| Missing (%) | 3.6 |

**Fig. S6 Summary of raw reads, read quantification and de novo assembly**

**A**, Summary of raw reads and read quantification with Salmon. **B**, Summary of the assembled transcriptome. **C**, Summary of transcriptome quality assessed by BUSCO.

**A**

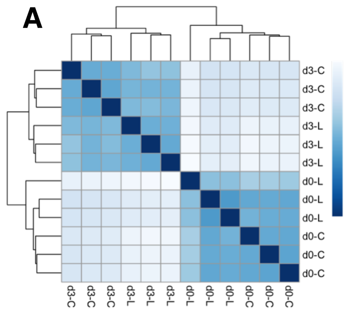

**B**

| Comparison between | | *padj* < 0.1 | *padj* < 0.05 | *padj* < 0.01 |
|---|---|---|---|---|
| Regions -- Lateral vs Central | Lateral | 1164 | 938 | 652 |
| | Central | 1165 | 866 | 469 |
| Time points -- 0 DPE vs 3 DPE | 0 DPE | 3870 | 3512 | 2839 |
| | 3 DPE | 3932 | 3531 | 2857 |

**C**

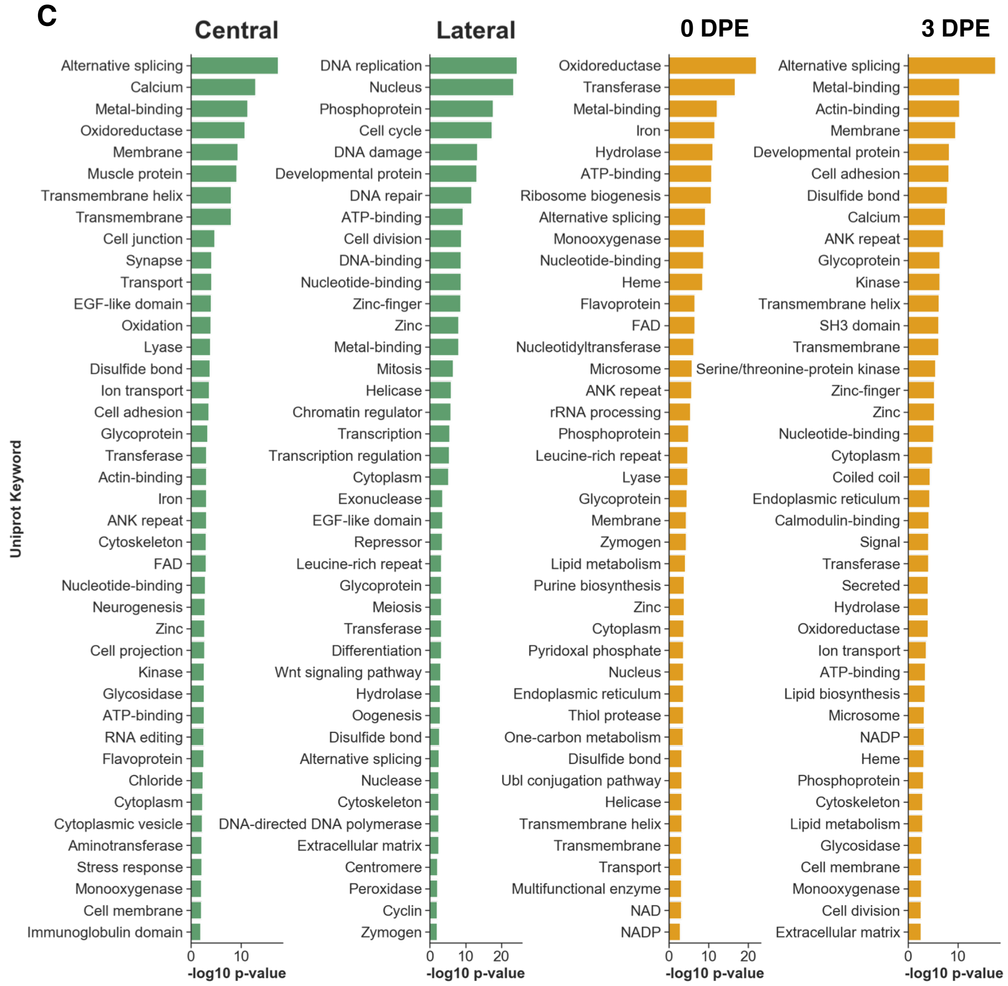

**Fig. S7 Summary of differentially expressed genes (DEG)**

**A**, Heatmap of sample-to-sample distance and hierarchical clustering. Color code indicates Euclidean distance. **B**, Number of upregulated DEG in each comparison. Numbers in different FDR (*padj*) cut-off are shown. **C**, Top 40 statistically enriched uniport keyword in transcripts upregulated in each group according to DAVID.

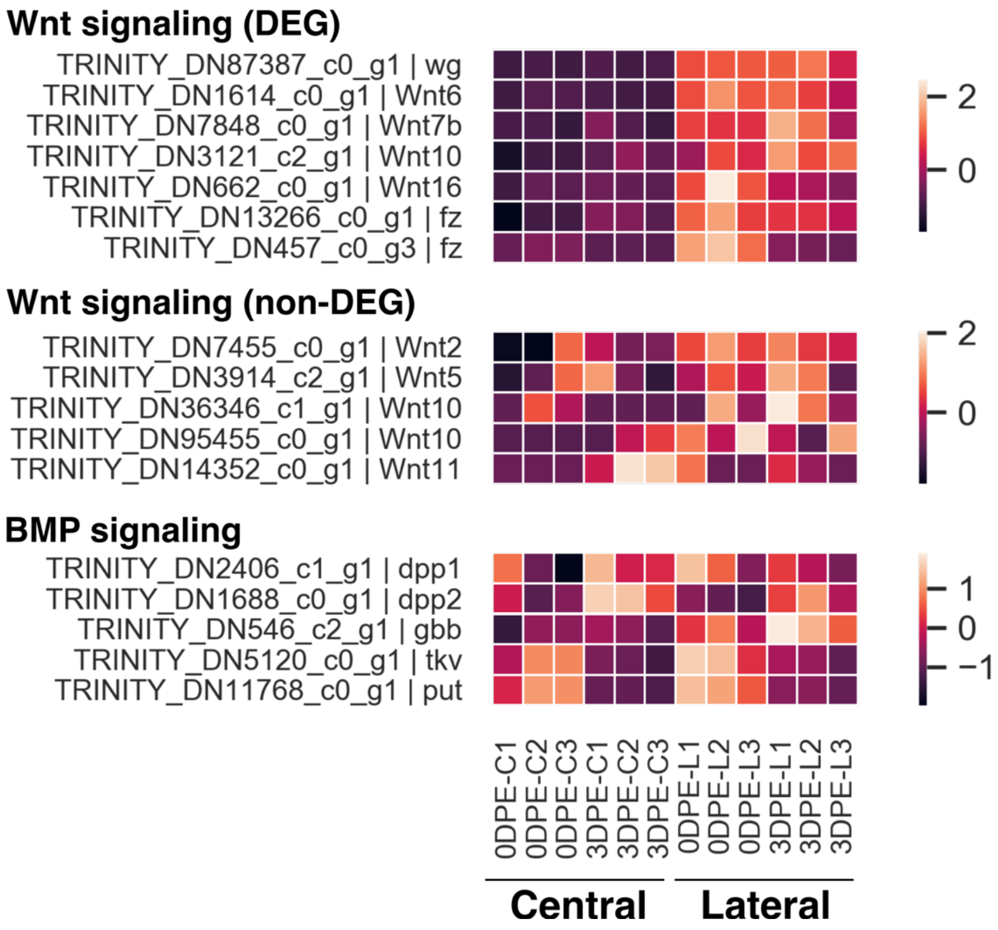

**Fig. S8 Expression profile of components of Wnt and BMP signaling**

Color code indicates z-score normalized TPM.

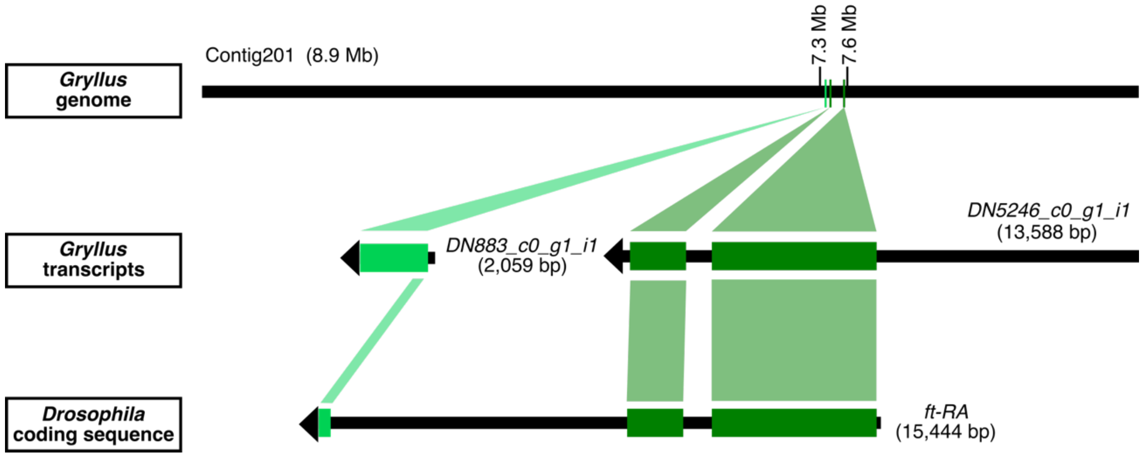

**Fig. S9 Two *ft* transcripts of *Gryllus* derive from a single genomic locus**

The contig201 in the *Gryllus* genome, and two transcripts of *ft* in the *Gryllus* transcriptome and the protein-coding sequence of a *ft* transcript in *Drosophila* are drawn with a black bar and black arrows, respectively. Color boxes indicate the corresponding regions among these nucleotide sequences according to a BLAST analysis. Two *ft* transcripts of *Gryllus* hit to the single *Drosophila* *ft* gene, and to the genomic locus within 300 kb. The order of the blast-hit sites on the genome agrees that two transcripts derive from a single *ft* gene.

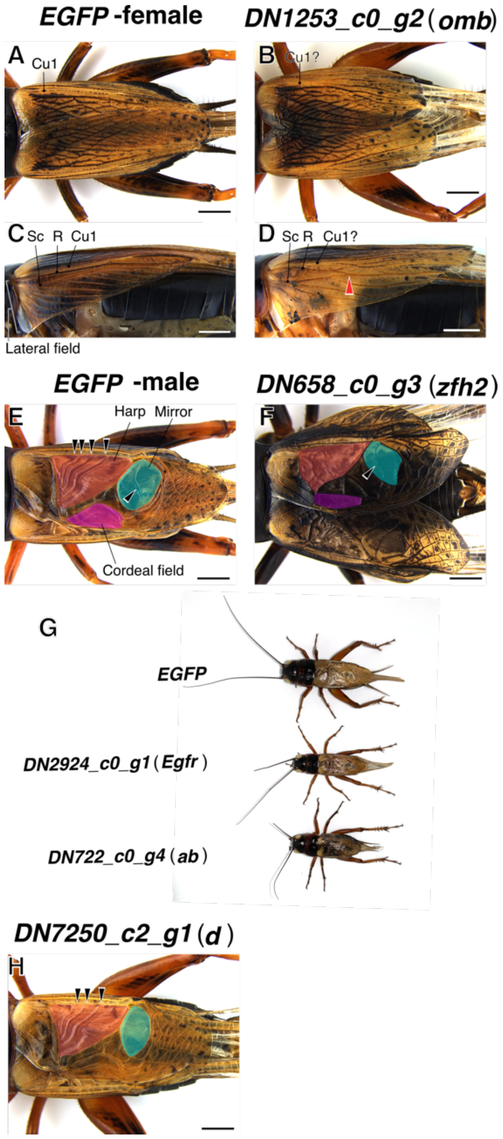

**Fig. S10 RNAi phenotypes of *omb*, *zfh2*, *Egfr* and *ab***

**A**–**D**, Dorsal (**A**, **B**) and lateral (**C**, **D**) views of female adults after *EGFP* (**A**, **C**) and *omb* (**B**, **D**) dsRNA injections. Characteristic veins were annotated following Schaffner & Koch (1987). Cu1, cubitus 1; R, radius; Sc, subcostal. R and Sc are fused (arrowhead) in the *omb* RNAi cricket but not in control. **E**, **F**, Dorsal views of male adults after *EGFP* (**E**) and *zfh2* (**F**) dsRNA injection. The *zfh2* RNAi cricket displays abnormal wing vein patterns represented as a loss of organized cross-veins in both the harp (red) and cordeal fields (magenta), and the deformed mirror (cyan). **G**, Adult male crickets after *EGFP*, *Egfr* and *ab* dsRNA injection at the same magnification. **H**, Dorsal view of adult males after *d* dsRNA injection showing loss of cross-veins in the harp and mirror. Mirror also shows a deformed shape compared to control (**E**). Scale bar is 2 mm.

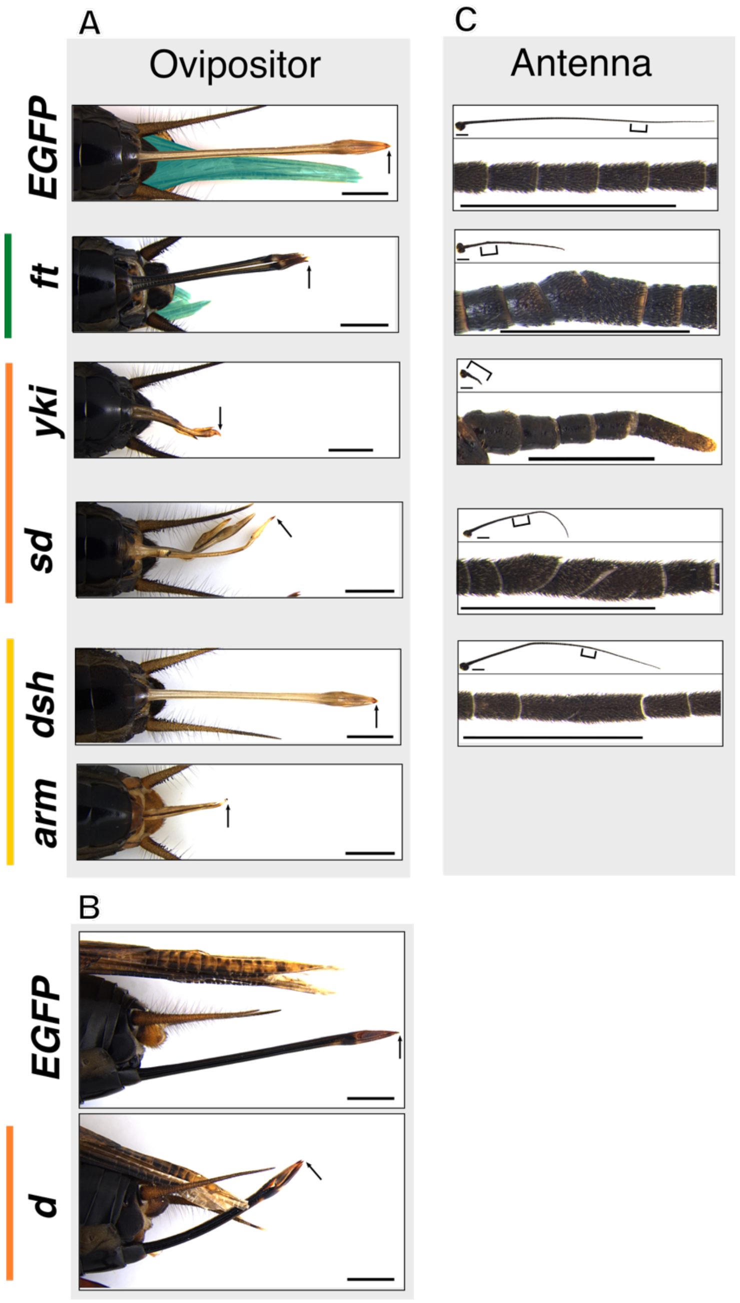

**Fig. S11 RNAi effects of Wnt/Ft-Ds/Hippo pathway components on ovipositors and antennae**

**A**, Ventral view of posterior-end regions of RNAi adult crickets. The hindwing is shaded in cyan. Arrow indicates a distal end of an ovipositor. Scale bar is 1 mm. **B**, Lateral views of posterior-end regions of *EGFP* and *d* RNAi crickets. **C**, Bracketed part of the whole antenna (top) is magnified in the bottom panel. Scale bars are 1 mm.

**Table S1 Summary of mosaic knockouts**

| sgRNA | No. of injected egg | No. of hatched nymph (%) | No. of embryo died just before hatching (%) | Phenotype (%) | | |
|---|---|---|---|---|---|---|
| | | | | No effect | Moderate | Strong |
| *EGFP* | 213 | 65 (30.5) | 2 (0.93) | 67 (100) | 0 (0) | 0 (0) |
| *apAB* | 215 | 19 (8.84) | 102 (47.4) | 5 (4.13) | 37 (30.6) | 79 (65.3) |
| *vg* | 212 | 45 (21.2) | 18 (8.49) | 38 (60.3) | 24 (38.1) | 1 (1.59) |

**Table S2 Summary of nRNAi-mediated functional screening**

| Target | FlyBase FBgn | Gene symbol | ug injected dsRNA / nymph | No. injected | No. of male adult | No. of female adult | %survival rate | Adult phenotype | | | | Note |
|---|---|---|---|---|---|---|---|---|---|---|---|---|
| | | | | | | | | Small wing size | Other wing phenotype | Other specific phenotype | wildtype | |
| EGFP | - | - | 5 | 13 | 8 | 4 | 92.3 | 0 | 0 | 0 | 12 | |
| TRINITY_DN1122_c0_g3 | FBgn0005771 | noc | 6.28 | 13 | 2 | 0 | 15.4 | 0 | 0 | 0 | 2 | |
| TRINITY_DN1253_c0_g2 | FBgn0000179 | bi (omb) | 6.69 | 13 | 3 | 3 | 46.2 | 0 | 6 | 0 | 0 | |
| TRINITY_DN13266_c0_g1 | FBgn0001085 | fz | 7.60 | 14 | 2 | 8 | 71.4 | 0 | 0 | 0 | 10 | |
| TRINITY_DN14433_c0_g1 | FBgn0037384 | dgrn | 7.33 | 12 | 10 | 1 | 91.7 | 0 | 0 | 0 | 11 | |
| TRINITY_DN1533_c0_g1 | FBgn0003892 | ptc | 6.75 | 13 | 4 | 6 | 76.9 | 0 | 0 | 0 | 10 | |
| TRINITY_DN1540_c2_g1 | No BLAST hit | Smbt2 | 6.79 | 13 | 7 | 3 | 76.9 | 0 | 0 | 0 | 10 | |
| TRINITY_DN1614_c0_g1 | FBgn0031902 | Wnt6 | 6.45 | 13 | 5 | 4 | 69.2 | 0 | 0 | 0 | 9 | |
| TRINITY_DN177_c0_g3 | FBgn0001075 | ft | 5.38 | 8 | 3 | 1 | 50.0 | 0 | 0 | 0 | 4 | |
| TRINITY_DN2297_c0_g2 | FBgn0000658 | fj | 7.14 | 12 | 7 | 2 | 75.0 | 0 | 0 | 0 | 9 | |
| TRINITY_DN263_c0_g2 | FBgn0263118 | tx | 6.09 | 14 | 6 | 5 | 78.6 | 0 | 0 | 0 | 11 | |
| TRINITY_DN2910_c3_g1 | FBgn0004583 | ex | 6.71 | 14 | 5 | 5 | 71.4 | 0 | 0 | 0 | 10 | |
| TRINITY_DN2924_c0_g1 | FBgn0003731 | Egfr | 7.09 | 12 | 3 | 4 | 58.3 | 0 | 0 | 7 | 0 | 7x Small body size |
| TRINITY_DN3121_c2_g1 | FBgn0031903 | Wnt10 | 7.43 | 13 | 3 | 5 | 61.5 | 0 | 0 | 0 | 8 | |
| TRINITY_DN3526_c3_g1 | FBgn0085424 | nub | 7.25 | 13 | 0 | 2 | 15.4 | 0 | 0 | 0 | 2 | |
| TRINITY_DN411_c6_g1 | FBgn0036494 | Toll-6 | 6.58 | 9 | 1 | 2 | 33.3 | 0 | 0 | 0 | 3 | |
| TRINITY_DN411_c6_g2 | FBgn0029114 | Tollo | 6.83 | 11 | 5 | 5 | 90.9 | 0 | 0 | 0 | 10 | |
| TRINITY_DN457_c0_g3 | FBgn0001085 | fz | 5.41 | 13 | 6 | 4 | 76.9 | 0 | 0 | 0 | 10 | |
| TRINITY_DN5246_c0_g1 | FBgn0001075 | ft | 6.26 | 13 | 8 | 0 | 61.5 | 7 | 0 | 0 | 1 | |
| TRINITY_DN5740_c0_g1 | FBgn0044028 | Notum | 7.22 | 14 | 7 | 1 | 57.1 | 0 | 0 | 0 | 8 | |
| TRINITY_DN5792_c0_g3 | FBgn0003450 | snk | 7.33 | 14 | 6 | 1 | 50.0 | 0 | 0 | 0 | 7 | |
| TRINITY_DN658_c0_g3 | FBgn0004607 | zfh2 | 6.08 | 13 | 3 | 0 | 23.1 | 0 | 3 | 0 | 0 | 3x irregular vein pattern; 2x failed to eclose completely |
| TRINITY_DN662_c0_g1 | FBgn0004360 | Wnt16 | 7.43 | 14 | 4 | 3 | 50.0 | 0 | 0 | 0 | 7 | |
| TRINITY_DN695_c0_g2 | FBgn0004839 | otk | 6.14 | 14 | 2 | 5 | 50.0 | 0 | 0 | 0 | 7 | |
| TRINITY_DN722_c0_g4 | FBgn0264442 | ab | 6.16 | 13 | 8 | 2 | 76.9 | 0 | 0 | 8 | 2 | 8x small body size |
| TRINITY_DN7250_c2_g1 | FBgn0262029 | d | 5.76 | 14 | 8 | 4 | 85.7 | 0 | 6 | 4 | 2 | 6x defects in male vein pattern (the AP vein in the mirror); 4x short and curved ovipositors |
| TRINITY_DN7341_c0_g2 | FBgn0000492 | Dr | 6.36 | 13 | 4 | 2 | 46.2 | 0 | 0 | 0 | 6 | |
| TRINITY_DN7377_c0_g1 | FBgn0004644 | hh | 6.38 | 14 | 4 | 5 | 64.3 | 0 | 0 | 0 | 9 | |
| TRINITY_DN7848_c0_g1 | FBgn0004360 | Wnt2 | 5.28 | 14 | 4 | 3 | 50.0 | 0 | 0 | 0 | 7 | |
| TRINITY_DN8291_c0_g1 | FBgn0039286 | dan | 1.54 | 14 | 6 | 4 | 71.4 | 0 | 0 | 0 | 10 | |
| TRINITY_DN87387_c0_g1 | FBgn0284084 | wg | 5.79 | 14 | 6 | 1 | 50.0 | 0 | 0 | 0 | 7 | |
| TRINITY_DN876_c5_g2 | FBgn0034476 | Toll-7 | 5.10 | 14 | 4 | 6 | 71.4 | 0 | 0 | 0 | 10 | |
| TRINITY_DN883_c0_g1 | FBgn0001075 | ft | 5.52 | 14 | 6 | 6 | 85.7 | 12 | 0 | 0 | 0 | |
| TRINITY_DN9479_c0_g1 | FBgn0052405 | Cpr65Av | 5.32 | 14 | 6 | 2 | 57.1 | 0 | 0 | 0 | 8 | |

**Table S3 Summary of additional nRNAi analysis**

| Target | Amount of injected dsRNA / nymph | Injected instar | No. injected | No. of male adult | No. of female adult | %survival rate | No. of small winged adult (%) | Note |
|---|---|---|---|---|---|---|---|---|
| Experiment #1 | | | | | | | | |
| *EGFP* | 3 μg | 3rd | 34 | 10 | 15 | 73.5 | 0 (0) | |
| *disheveled* (*dsh*) | 3 μg | 3rd | 29 | 10 | 13 | 79.3 | 23 (100) | |
| *yorkie* (*yki*) | 3 ng | 3rd | 6 | 0 | 0 | 0.0 | *N.A.* | |
| *scalloped* (*sd*) | 3 μg | 3rd | 35 | 10 | 11 | 60.0 | 21 (100) | Other phenotype: 21 x short antennae; 11 x short ovipositors |
| Experiment #2 | | | | | | | | |
| *EGFP* | 50 ng | 6th | 23 | 5 | 7 | 52.2 | 0 (0) | |
| *yorkie* (*yki*) | 30 ng | 6th | 22 | 0 | 0 | 0.0 | *N.A.* | |
| *yorkie* (*yki*) | 3 ng | 6th | 24 | 8 | 8 | 66.7 | 16 (100) | Other phenotype: 16 x short antennae and ovipositors |
| Experiment #3 | | | | | | | | |
| *EGFP* | 5 μg | 3rd | 15 | 8 | 3 | 73.3 | 0 (0) | |
| *armadillo* (*arm*) | 10 ng | 3rd | 17 | 0 | 0 | 0.0 | *N.A.* | |
| *armadillo* (*arm*) | 1 ng | 3rd | 18 | 2 | 2 | 22.2 | 4 (10) | Other phenotype: 2 x short ovipositor, 3 x short antennae, 1 x small body size |
| Experiment #4 | | | | | | | | |
| *EGFP* | 5 μg | 3rd | 13 | 0 | 7 | 53.8 | 0 (0) | |
| *armadillo* (*arm*) | 1 ng | 3rd | 24 | 3 | 6 | 37.5 | 8 (88.8) | Other phenotype: 8 x short antennae and ovipositors and small body size |

**Table S4 Oligonucleotides used**

| Target | Purpose | Sequence #1 | Sequence #2 | bp product size |
|---|---|---|---|---|
| *in situ* hybridization | | | | |
| *wingless* (*wg*) | PCR primer | atttaggtgacactatagaaCTCTGCGGGAGAAGATGAAC | taatacgactcactatagggCACGTCTTGCACTTGACCTC | 523 |
| *apterous AB* (*apAB*) | PCR primer | atttaggtgacactatagaaATCTGAACGCGGACTACCTG | taatacgactcactataggGTCTTCTGCGAGAGCTGCTT | 223 |
| *vestigial* (*vg*) | PCR primer | atttaggtgacactatagaaCAGTACGTCTCCGCCAACTG | taatacgactcactataggGCCTGGTAGTTGCTGTTCCA | 188 |
| Generation of *vg* reporter line | | | | |
| Oligonucleotides for sgRNA synthesis | | | | |
| *vg* 5′ region | sgRNA template | taatacgactcactatagACTGATCGCCCTCGA | ttctagctctaaaacGGCACTCGAGGGCGATCAG | *N.A.* |
| G'act-eGFP donor plasmid | sgRNA template | taatacgactcactatagGATGTCCCAGGCGAA | ttctagctctaaaacGCCCTTCGCCTGGGACATC | *N.A.* |
| Primers for genomic PCR | | | | |
| Junction sequence between genome and the donor | PCR primer | TTCCTCGACGAGTGTGTCTG | GCTTCGAAAAGTGTCGGCTA | 354 |
| Expression of ecdysone responsive genes | | | | |
| E74 | qPCR primer | ATGCTCGACCTGGGCTTCCA | AGAACCTTCGCGGCTCTTGC | 108 |
| E75 | qPCR primer | CGCCTCGCCTGTATGTTCGA | GCGTTCGACGACGAGTGGAT | 95 |
| *Ribosomal protein L32* (*RpL32*) | qPCR primer | GATTCGCCCAGTTTATCGTC | GGCTTCAGCTTCTTGTACCG | 90 |
| nRNAi-mediated functional screening | | | | |
| TRINITY_DN1122_c0_g3 (noc) | PCR primer | taatacgactcactataggGACGACTTGGAGGGCTTCTC | taatacgactcactataggGAACACTGAACCGGGAGTGG | 131 |
| TRINITY_DN1253_c0_g2 (bi) | PCR primer | taatacgactcactataggGCGAGTCAGGGTGGATGTAC | taatacgactcactatagggTCCCGCAGATGAAATTCCGT | 204 |
| TRINITY_DN14433_c0_g1 (dgrn) | PCR primer | taatacgactcactatagggTTTCCTAGCACGATGGCTCC | taatacgactcactatagggCTGAGGGACTGTTGAGGTGG | 200 |
| TRINITY_DN1533_c0_g1 (ptc) | PCR primer | taatacgactcactataggGAATTGGTGGTGGCTGCAAG | taatacgactcactatagggACATGTGCATGGCAGGAAGA | 199 |
| TRINITY_DN1540_c2_g1 (Smbt2) | PCR primer | taatacgactcactatagggACGGCTGGCAGTGATTAAGT | taatacgactcactatagggTGGCTTCACTGCTACCCAAG | 203 |
| TRINITY_DN1614_c0_g1 (Wnt6) | PCR primer | taatacgactcactatagggTCGACCTGTCTTGTGCACTC | taatacgactcactataggGCAAGGTCAGCGTCAAAGTG | 200 |
| TRINITY_DN177_c0_g3 (ft) | PCR primer | taatacgactcactataggGAGATCGAGCGCCTCAACTC | taatacgactcactataggGAGCTGTGGTTGTCCAGGTG | 215 |
| TRINITY_DN2297_c0_g2 (fj) | PCR primer | taatacgactcactataggGAATCGCAATGAGCACGGTC | taatacgactcactataggGCGGCTGCATATAAATCGCC | 200 |
| TRINITY_DN263_c0_g2 (tx) | PCR primer | taatacgactcactataggGCTGTCACGAGAAGAGGTCC | taatacgactcactatagggATGTCTCCGCACGACTTGTT | 199 |
| TRINITY_DN2910_c3_g1 (ex) | PCR primer | taatacgactcactatagggAGCATTGTACGGTGGTGGAG | taatacgactcactataggGCGCAACCATTAGCACTACG | 200 |
| TRINITY_DN2924_c0_g1 (Egfr) | PCR primer | taatacgactcactatagggTTCACAATGCCAGAACGGGA | taatacgactcactatagggTGTTTAGGACCACGACAGCC | 197 |
| TRINITY_DN3121_c2_g1 (Wnt10) | PCR primer | taatacgactcactatagggTGCCATCGAGGAGTAGGTCA | taatacgactcactatagggCATCTTCTGCCGCATTGACG | 209 |
| TRINITY_DN3526_c3_g1 (nub) | PCR primer | taatacgactcactatagggCCCAAGAACAGGCTGAGGAG | taatacgactcactataggGCAAAGTCACAACAGCCCTG | 126 |
| TRINITY_DN411_c6_g1 (Toll-6) | PCR primer | taatacgactcactatagggTCGCACTCGACAAACAGTTC | taatacgactcactatagggCTTCAACCACTTCGTGCTGA | 181 |
| TRINITY_DN411_c6_g2 (Tollo) | PCR primer | taatacgactcactatagggCAGTTGGTAGGGCACGTCAT | taatacgactcactatagggAGTGCTCGAAATCAGCCGAA | 200 |
| TRINITY_DN457_c0_g3 (fz) | PCR primer | taatacgactcactatagggAGACTTCGCTAAGGCACCTG | taatacgactcactataggGCCACGAATCCTTTCTGTGC | 199 |
| TRINITY_DN5246_c0_g1 (ft) | PCR primer | taatacgactcactatagggAACGACAACCCGCCCATATT | taatacgactcactatagggCGGCTCGGTGGTAGAAATGT | 201 |
| TRINITY_DN5740_c0_g1 (Notum) | PCR primer | taatacgactcactatagggAGACGAGCTACGCGAGAGTA | taatacgactcactatagggCTTTGGTTGCACGCCCAAAT | 223 |
| TRINITY_DN5792_c0_g3 (snk) | PCR primer | taatacgactcactataggGACTTCAGTCGGCCTTGTGA | taatacgactcactatagGGGATGGTACTTTCACTGCCA | 195 |
| TRINITY_DN658_c0_g3 (zfh2) | PCR primer | taatacgactcactatagggTGCGCTCTGTACTGCATCAA | taatacgactcactatagggTGCCACAACTACTGTTCGCT | 201 |
| TRINITY_DN662_c0_g1 (Wnt16) | PCR primer | taatacgactcactatagggTGGATGTGCACTAGATCGCC | taatacgactcactatagggTCGATGCCATGGAATCTCCG | 198 |
| TRINITY_DN695_c0_g2 (otk) | PCR primer | taatacgactcactatagggAAGACGGTGATCCTGGAGGT | taatacgactcactatagggCCACCAACTCCTCCAGGAAC | 239 |
| TRINITY_DN722_c0_g4 (ab) | PCR primer | taatacgactcactataggGGAAGGCACTCCCTCACTTC | taatacgactcactatagGGGTGATTGTCGTCCATCGT | 201 |
| TRINITY_DN7250_c2_g1 (d) | PCR primer | taatacgactcactatagggCGGCAATACAGAGCTTTCGC | taatacgactcactatagggTGGTCAGGGACTAGAGGTGG | 201 |
| TRINITY_DN7341_c0_g2 (Dr) | PCR primer | taatacgactcactataggGCAAGAGTCATACACGCACG | taatacgactcactatagggTGCTTCGGAAACAATGTGCG | 200 |
| TRINITY_DN7377_c0_g1 (hh) | PCR primer | taatacgactcactatagggATGCTCTCTCTCACTGTGCG | taatacgactcactatagggAGGCCCAGCTTTGAACACTT | 200 |
| TRINITY_DN7848_c0_g1 (Wnt2) | PCR primer | taatacgactcactataggGCATGCTCACACACATCCAC | taatacgactcactatagggAGTCGAGCAATACACGTGCA | 176 |
| TRINITY_DN8291_c0_g1 (dan) | PCR primer | taatacgactcactatagggACCTGGTCTACGTGCCATTG | taatacgactcactatagggAGCTCTGCACAGACAATGCT | 122 |
| TRINITY_DN87387_c0_g1 (wg) | PCR primer | taatacgactcactataggGAAGCACTGTACGGCTGGAA | taatacgactcactatagggAGTTCACTGTCCGCCGAAAT | 196 |
| TRINITY_DN876_c5_g2 (Toll-7) | PCR primer | taatacgactcactatagggAGCTTCTCGTGCTCGATCTG | taatacgactcactataggGTCCTCGATGTAGCCGATGG | 200 |
| TRINITY_DN883_c0_g1 (ft) | PCR primer | taatacgactcactatagggACAAGCAACCAGTCCTCAGG | taatacgactcactatagggCCAAGCCCTCCGAAGAGTAC | 191 |
| TRINITY_DN9479_c0_g1 (Cpr65Av) | PCR primer | taatacgactcactatagggCCCCGTTTTCGGGTATAAAT | taatacgactcactatagggAGAGCTGCCGAGGAGATGT | 125 |
| Additional nRNAi | | | | |
| *disheveled* (*dsh*) | PCR primer | taatacgactcactatagggCACGCACATCTTCCTTCTCA | taatacgactcactatagGGCTCTATGCGACCATCAAT | 196 |
| *armadillo* (*arm*) | PCR primer | taatacgactcactatagggATCCAAGTCAAAGGCTGGTG | taatacgactcactataggGCGCTGATTGTTGCAAGTTA | 185 |
| *yorkie* (*yki*) | PCR primer | taatacgactcactatagggCCCTGCAGATGGAGAGAGAG | taatacgactcactatagggTGATCAGTCACGCCTGAGA | 143 |
| *scalloped* (*sd*) | PCR primer | taatacgactcactatagggCCAGGTGTTGGCTAGAAGGA | taatacgactcactatagggCCTGGGTAGGACACTGGAGA | 207 |

Small letters indicate universal sequences
